# Control of cholesterol-induced adipocyte inflammation by the Nfe2l1-Atf3 pathway

**DOI:** 10.1101/2024.07.22.604614

**Authors:** Carolin Jethwa, Anne Hoffmann, Stefan Kotschi, Janina Caesar, Matthias Kern, Anna Worthmann, Christian Schlein, Sajjad Khani, Bernardo A. Arús, Oliver T. Bruns, Adhideb Ghosh, Christian Wolfrum, Yvonne Döring, Stephan Herzig, Christian Weber, Matthias Blüher, Scott B. Widenmaier, Gökhan S. Hotamışlıgil, Alexander Bartelt

## Abstract

While adipocytes are critical pillars of energy metabolism, their dysfunction is linked to adipose tissue (AT) inflammation, insulin resistance, and ectopic lipotoxicity in cardiometabolic diseases. However, the mechanisms causing adipocyte inflammation and insulin resistance remain unclear. Here, we show that excess cholesterol induces adipocyte dysfunction, which is suppressed by the transcription factor Nfe2l1 (nuclear factor erythroid derived-2, like-1). Nfe2l1 is required to sustain proteasome function in adipocytes and proteotoxic stress induces adipocyte inflammation via the activation of Atf3. In humans, the Nfe2l1-proteasome pathway is inversely correlated to body mass index (BMI) in an adipose-depot specific manner. In mice, loss of adipocyte Nfe2l1 caused AT inflammation with a pronounced infiltration of macrophages and T cells. Mice lacking adipocyte Nfe2l1 displayed severe adipocyte dysfunction during diet-induced obesity (DIO), characterized by lower adipokine levels, steatosis, glucose intolerance and insulin resistance. *Nfe2l1*^ΔAT^ mice on an Apoe-deficient (*Apoe*^−/−^) background fed a cholesterol-rich Western Diet (WD), developed a lipoatrophy-like syndrome, dyslipidemia, and enhanced atherosclerosis. Our results reveal an important role for proteasome-mediated proteostasis in adipocytes and indicate that Nfe2l1 is linked to metabolic health in humans and preclinical mouse models. Promoting proteostasis in adipocytes may thus alleviate inflammation in obesity, potentially averting adverse cardiometabolic outcomes.

**Graphical abstract:** 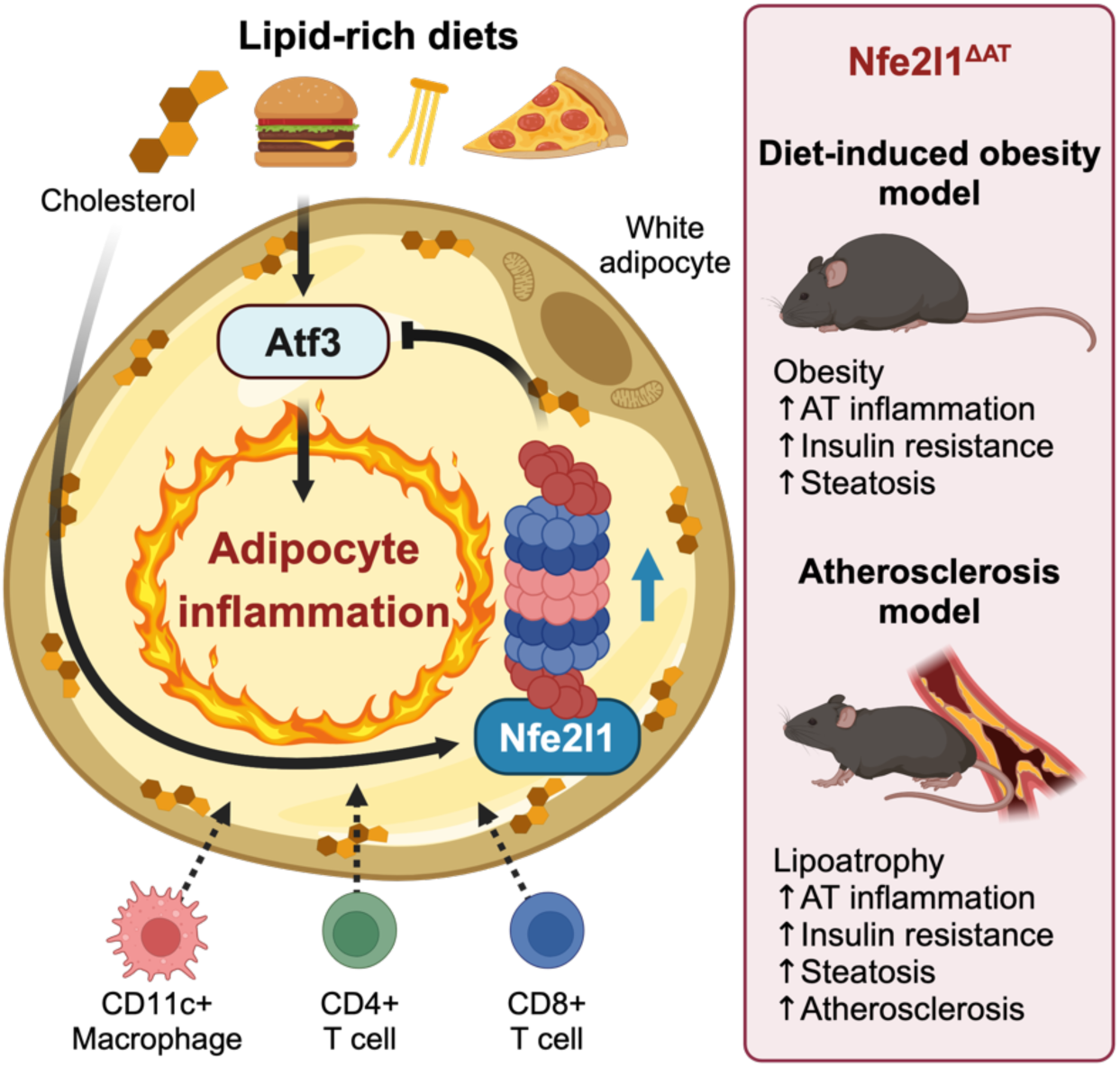

## Introduction

Obesity is characterized by progressive adipocyte dysfunction, which is associated with increased risk of developing type 2 diabetes (T2D), cardiovascular disease (CVD), and other comorbitities^1–4^. Conversely, the relative absence of healthy adipocytes, as observed in congenital or acquired lipodystrophy, also increases cardiometabolic risk, mirroring the metabolic disturbances seen in obesity^5–7^. The inability to respond to insulin and other hormonal cues results in diminished lipid storage and elevated release of non-esterified fatty acids (NEFAs) into the circulation, leading to ectopic lipid accumulation. The state of insulin resistance is associated with aberrant metabolism of lipids, glucose, and proteins all of which are sensed at the endoplasmic reticulum (ER). The ER is a highly adaptive organelle, which orchestrates the activation of stress resistance pathways to maintain cellular homeostasis^8,9^. However, in obesity, chronic ER stress and inflammation become maladaptive and contribute to the progression of metabolic diseases^10^. In obesity, adipocytes display increased demand for protein synthesis during expansion and secretion of adipokines^11^. Moreover excessive nutrients such as cholesterol and NEFAs, as well as pro-inflammatory cytokines^11^ disrupt ER homeostasis, which also is linked to impaired proteostasis. Especially, excessive cholesterol, which accumulates in adipocytes in its free form during obesity, is toxic since it decreases membrane fluidity, alters membrane protein function, and ultimately leads to cellular dysfunction. The ER-resident transcription factor Nfe2l1 has been shown to regulate both proteostasis and cholesterol homeostasis in hepatocytes. Nfe2l1 is a cholesterol sensor and turnover, post-translational processing, and activity of Nfe2l1 are modulated by cholesterol^12^. In the liver, Nfe2l1 protects from hepatic steatosis and inflammation by maintaining proteostasis and inducing cholesterol removal^13,14^. Additionally, in cardiomyocytes, Nfe2l1 promotes heart regeneration following myocardial infarction^15^. We demonstrated that Nfe2l1 is essential for the adaptation of brown adipose tissue (BAT) to cold^16^. The absence of Nfe2l1 in brown adipocytes is linked to the whitening of BAT and diminished non-shivering thermogenesis. However, few studies have investigated the role of Nfe2l1 in white adipocytes. Similar to what was observed in thermogenic adipocytes^16^, mice lacking white adipocyte Nfe2l1 display white adipocyte abnormalities and altered responses to adrenergic stimulation or rosiglitazone treatment^17–19^. We have previously shown that in isolated adipocytes, proteasome function is important for adipocyte function and linked to inflammation^20^ and ferroptotic cell death^21,22^. Regardless of these studies, the role of white adipocyte Nfe2l1 in obesity and its impact on cardiometabolic health remain largely elusive.

## Results

### The Nfe2l1-proteasome-pathway is inversely correlated with human obesity

Nfe2l1 undergoes complex posttranslational processing, inducing the transcription of proteasomal subunit genes (Fig. 1A,B) when proteasomal activity is insufficient for cellular needs^14,23,24^, but the role of Nfe2l1 in white adipocytes remains largely unclear. Therefore, we investigated the gene expression of the Nfe2l1-proteasome pathway in samples obtained from human subjects (n=478) with a broad range of BMI by RNA sequencing (RNA-Seq). We found a positive correlation between *NFE2L1* mRNA expression and most proteasomal subunit (*PSM*) genes in subcutaneous white adipose tissue (SCAT) and visceral white adipose tissue (VISAT), highlighting the strong link between *NFE2L1* and the proteasome in human adipose tissue (Fig. 1C). Moreover, we found that *NFE2L1* expression in both SCAT and VISAT was inversely correlated with BMI (Fig. 1C,D). This indicates that obesity is potentially associated with compromised Nfe2l1-mediated proteostasis in WAT.

**Fig. 1:**
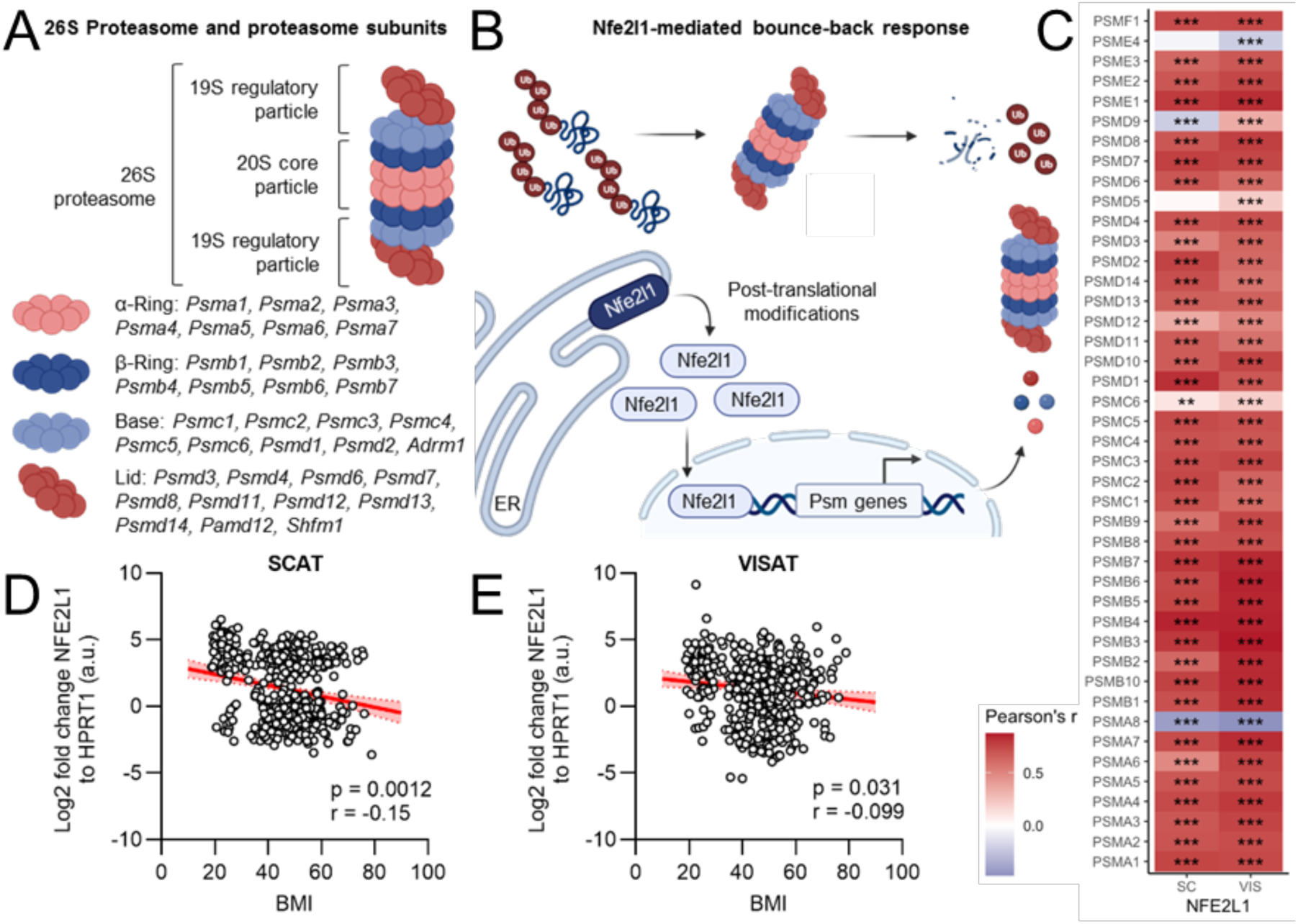
The Nfe2l1-proteasome pathway in adipose tissue is inversely correlated with human obesity. (A) Scheme of the 26S proteasome and proteasome subunits. (B) Model of Nfe2l1 action. ER: Endoplasmic reticulum, Psm genes: proteasome subunit genes, Ub: Ubiquitin. (C) Correlation of *NFE2l1* expression in SCAT and VISAT with expression of proteasomal subunit genes in the LOBB (n=1,479). \*\**P*<0.01, \*\*\**P*<0.001 by multiple-linear regression with Pearson’s correction. (D) Correlation of *NFE2L1* expression with BMI in SCAT of non-obese and obese human individuals in the LOBB (n=478). Data are mean + s.e.m, *P* by linear regression with Spearman’s correction. (E) Correlation of *NFE2L1* expression with BMI in VISAT of non-obese and obese human individuals in the LOBB (n=478). Data are mean+s.e.m, *P* by linear regression with Spearman’s correction.

### Nfe2l1-mediated proteostasis prevents inflammation in white adipocytes

It is unclear if the lower expression of NFE2L1 at higher BMI is specific to adipocytes and is causally involved in adipocyte proteostasis and the development of cardiometabolic diseases. To address these questions, we used a Nfe2l1 Cre-loxP model crossed with Adipoq-Cre and generated an adipocyte-specific Nfe2l1 knockout mouse (*Nfe2l1*^ΔAT^) model (Supplementary Fig. 1). To test the effect of loss of *Nfe2l1* on proteostasis and cellular health, we isolated primary white adipocytes of *Nfe2l1*^ΔAT^ and floxed control (*flox/flox*) mice. *Nfe2l1*^ΔAT^ adipocytes displayed lower expression of proteasome subunit genes at baseline, and failure to induce their expression when treated with the proteasome inhibitor epoxomicin (Fig. 2A) compared to control cells. This was accompanied by higher expression of the ER stress surrogate marker genes *Ddit3*, *Hspa5* and *Xbp1s* as well as the inflammation surrogate marker genes *Ccl2* and *Saa3* (Fig. 2B). Notably, treatment with Epoxomicin, a chemical proteasome inhibitor, amplified this inflammatory stress response in the absence of *Nfe2l1* (Fig. 2B). On a functional level, lower expression of proteasome subunit genes in *Nfe2l1*^ΔAT^ adipocytes indeed translated to lower activity of all three proteolytic domains of the proteasome (Fig. 2C). This was linked to activation of stress kinase signaling pathways including p38 MAPK, Nf-KB and elF2a (Fig. 2D-F).

**Fig. 2:**
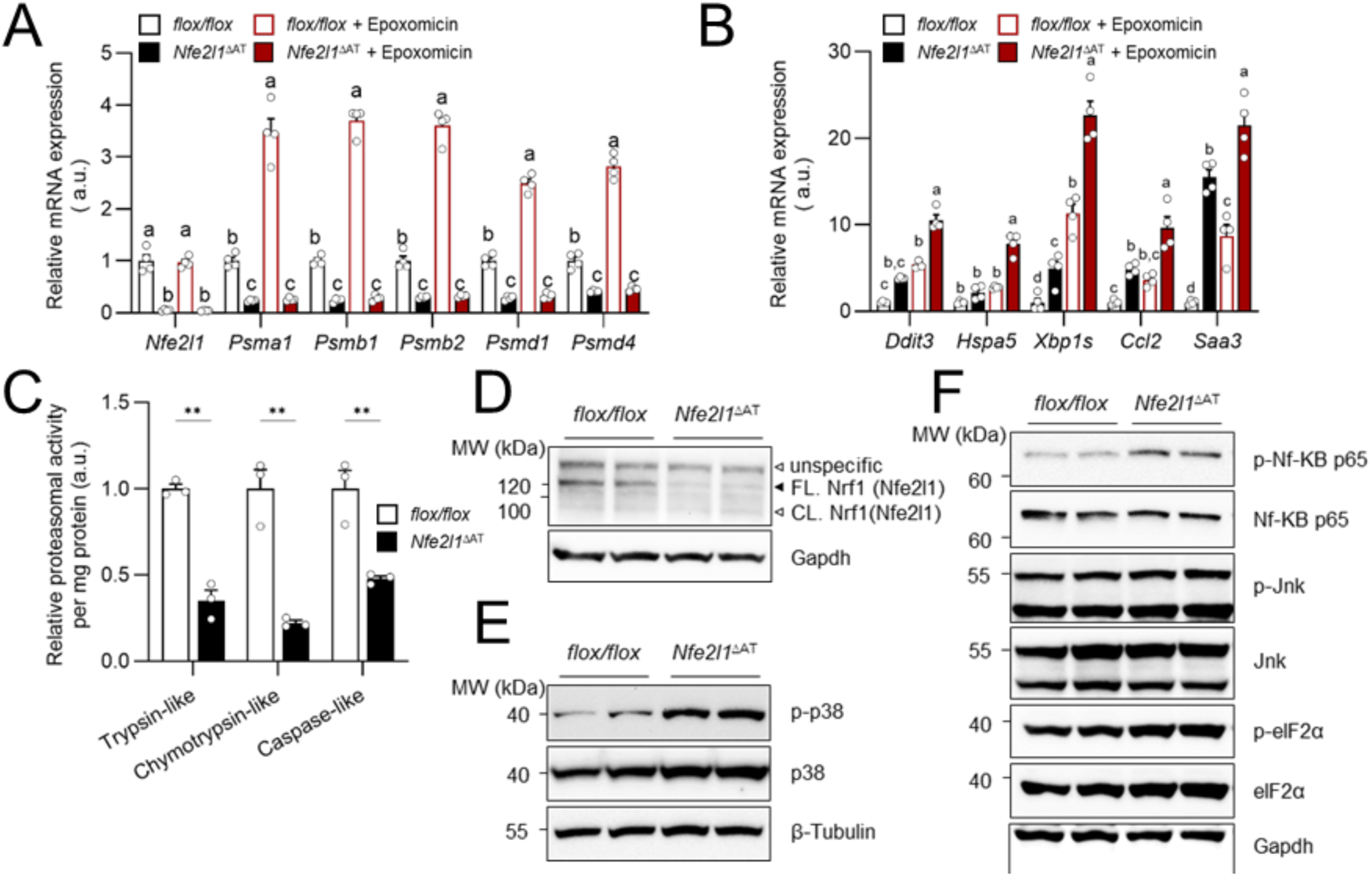
Impaired proteasome activity in Nfe2l1 knockout adipocytes is linked to inflammatory signaling. (A-B) Relative mRNA levels in primary white adipocytes of *flox/flox* and *Nfe2l1*^ΔAT^ mice, n=4, data are mean+s.e.m, different letters indicate significant differences (*P*_adj_<0.05) between groups tested by 1-way ANOVA with Tukey post-hoc test. (C) Proteasome activity in primary white adipocytes of *flox/flox* and *Nfe2l1*^ΔAT^ mice, n=3, data is mean+s.e.m, \*\**P*<0.01 by Student’s T-test. (D-E) Representative immunoblots of primary white adipocytes of *flox/flox* and *Nfe2l1*^ΔAT^ mice, n=2. FL.: full-length, CL.: cleaved.

### Atf3 mediates lipotoxicity-induced inflammation in white adipocytes

Considering the role of Nfe2l1 as a cholesterol sensor^12^, we hypothesized that the function of Nfe2l1 and proteostatic outcome might be linked to cholesterol homeostasis in adipocytes. To study this, we used methyl-beta-cyclodextrin (MbCD)-conjugated cholesterol, which results in ER cholesterol to mimic cholesterol overload of the cell. This treatment by itself, as well as in combination with proteasome inhibition by epoxomicin, led to a robust induction of inflammation surrogate marker genes *Ccl2* and *Cxcl10*. ER stress surrogate marker genes such as *Hspa5* and *Ddit3* were unaffected by any of the treatments. However, we observed a strong induction of the stress marker *Atf3* by epoxomicin treatment, which was amplified by the addition of cholesterol (Fig. 3A). Atf3 is member of the ATF/cAMP response element-binding family of transcription factors^25^ and a downstream target of Atf4 in the PERK pathway of the UPR^26^. The Atf3 promotor region contains multiple binding sites for other transcription factors, including NF-κB, indicating its induction by stress and inflammation signals^27^. Thus, we hypothesized that Atf3 might be involved in mediating the inflammatory response to cholesterol overload. Indeed, when Atf3 was silenced in 3T3-L1 adipocytes (Fig. 3B), this resulted in markedly lower expression of *Ccl2* in the cholesterol- and epoxomicin-treated group (Fig. 3C). However, this was independent of acute silencing of *Nfe2l1*. To achieve a better and physiological understanding of how cholesterol overload affects white adipocytes and how Nfe2l1 and Atf3 modulate this response, we performed RNA-Seq of primary white adipocytes transfected with *siNfe2l1* and *siAtf3* and treated with cholesterol (Fig. 3D). Notably, cholesterol treatment had a strong impact on the adipocyte transcriptome in all groups, whereas neither the silencing of *Atf3* nor the silencing of *Nfe2l1* by themselves had major effects (Fig. 3E). Thus, we decided to uncouple the effect of the treatment from the genetic manipulation to unravel how cholesterol treatment affected adipocytes. To start, we used gene enrichment analysis of the significantly up-regulated genes. The five most significantly enriched terms were “Response to unfolded protein”, “Regulation of DNA-templated transcription in response to stress”, “Response to lipids”, “Negative regulation of cell population proliferation” and “Inflammatory response”. This indicates that cholesterol overload induced a lipotoxic stress response. Notably, amongst the most up-regulated genes were genes coding for important inflammatory mediators such as *Cxcl1*, *Cxcl2* and *Cxcl10*, as well as stress marker genes such as *Hspa1a*, *Hspa1b*, *Ddit3* and *Atf3* (Fig. 3F).

**Fig. 3:**
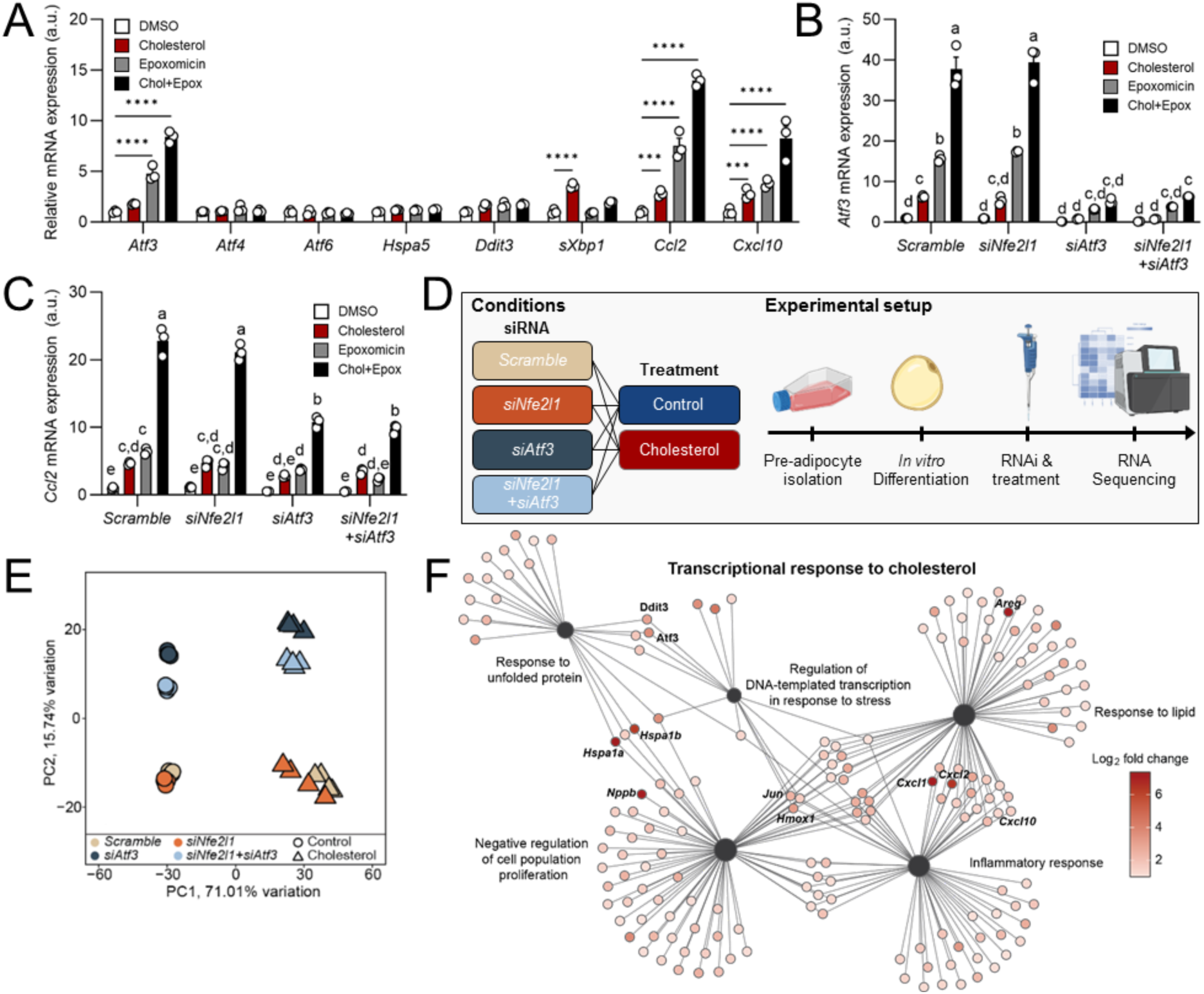
Cholesterol induces a lipotoxic stress response in white adipocytes. (A) Relative mRNA expression in 3T3-L1 adipocytes treated with 400 µM cholesterol, 100 nM epoxomicin or the combination of both (Chol+Epox) for 6 h, n=3, data are mean+s.e.m, \*\*\**P*_adj_<0.001, \*\*\*\**P*<0.001 by 1-way ANOVA with Dunnett‘s post hoc test. (B-C) Relative mRNA expression in 3T3-L1 adipocytes treated with 400 µM cholesterol, 100 nM epoxomicin or the combination of both for 6 h, n=3, data is mean+s.e.m, different letters indicate significant differences (*P*_adj_<0.05) between groups tested by 2-way ANOVA with Tukey post-hoc test. (D) Schematic workflow of RNA-Seq of primary white adipocytes treated with 400 µM cholesterol for 6 h. (E) Principal component analysis plot, n=4. (F) Top five up-regulated GO terms by cholesterol treatment (log2 fold change >1). Data is mean log2 fold change, n=4.

Gene enrichment analysis for the molecular function “DNA-binding transcription factor activity” demonstrated that Atf3 is the most highly induced transcription factor by cholesterol treatment in adipocytes (Fig. 4A). To understand how Atf3 affects the cholesterol-induced transcriptome, we investigated how silencing of *Atf3* affected the five most upregulated pathways and found that silencing of Atf3 significantly attenuated all of them (Fig. 4B). We also independently validated these findings in 3T3-L1 adipocytes, which showed activation of inflammatory and stress kinase signaling pathways NF-κB, Jnk and p38 with cholesterol treatment, as well as amplified signal by the addition of epoxomicin to the cholesterol treatment.

**Fig. 4:**
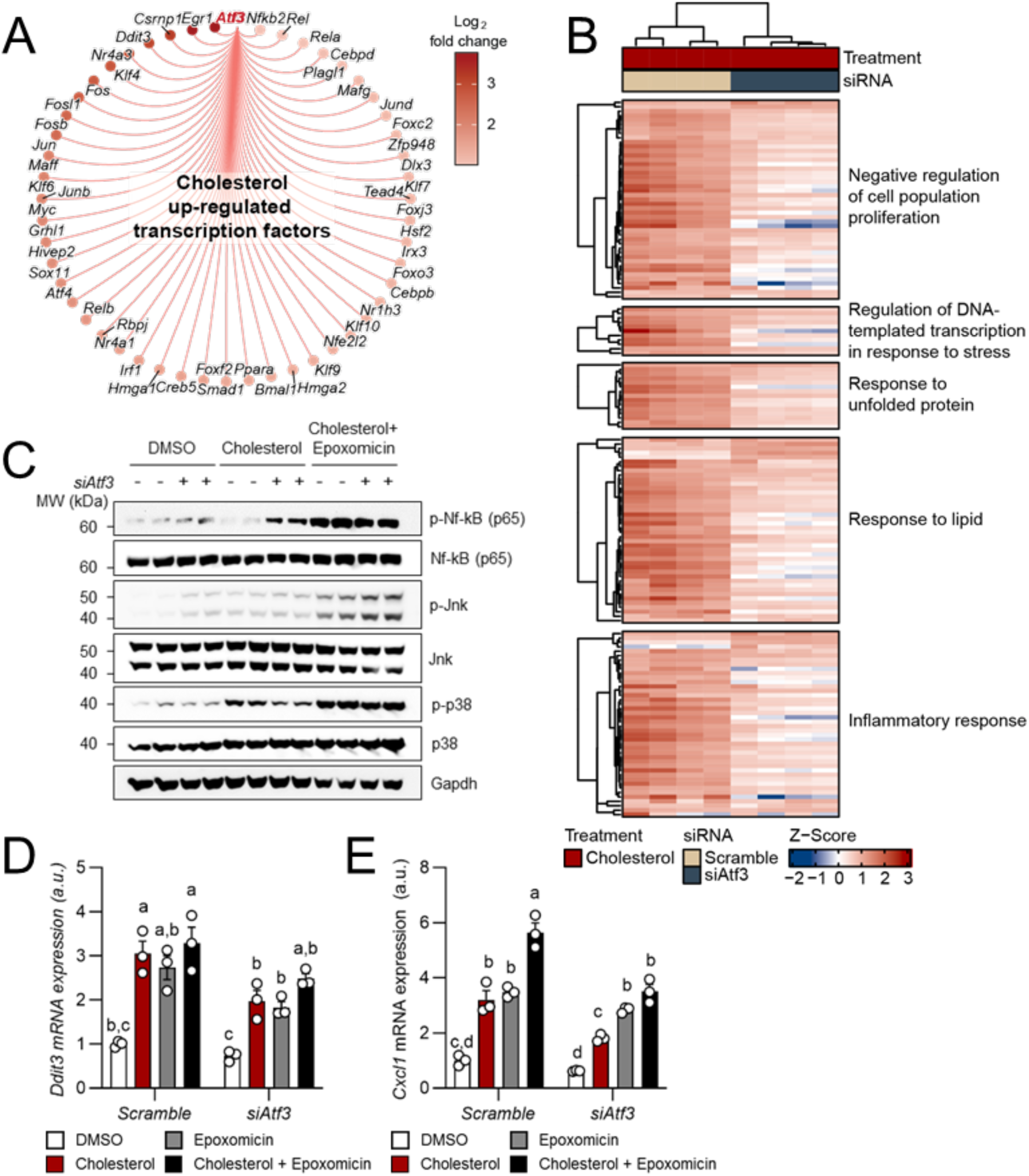
Atf3 mediates cholesterol-induced inflammation in white adipocytes. (A) Up-regulated transcription factors by cholesterol in primary white adipocytes, n=4. (B) Regulation of the top 5 cholesterol-induced GO terms by siAtf3 in primary white adipocytes, n=4. (C) Representative immunoblots of 3T3-L1 adipocytes transfected with siAtf3 and treated with cholesterol with or without epoxomicin for 6 h, n=2. (D-E) *Ddit3* and *Cxcl1* mRNA expression in 3T3-L1 adipocytes transfected with siAtf3 and treated with cholesterol with or without epoxomicin for 6 h, n=3, data are mean+s.e.m, different letters indicate significant differences (*P*_adj_<0.05) between groups tested by 2-way ANOVA with Tukey post-hoc test.

However, silencing of *Atf3* here did not affect activation of these pathways except for phosphorylation of p38 (Fig. 4C). On the gene expression level, silencing of *Atf3* was sufficient to lower *Ddit3* and *Cxcl1* expression in response to cholesterol treatment (Fig. 4D-E).

### Adipocyte Nfe2l1 protects cardiometabolic health in diet-induced obesity and atherosclerosis

Next, we investigated the role of adipocyte Nfe2l1 in vivo in the context of metabolic challenges in mice. On the tissue level, loss of adipocyte Nfe2l1 resulted in a complex adipose tissue inflammation with adipocyte hypertrophy, diminished browning and crown-like structures in both SCAT and GWAT, even in the absence high-fat diet (HFD) feeding (Fig. 5A). Moreover, RNA sequencing of SCAT, demonstrated that upregulated genes where enriched in pathways linked to immune cell infiltration and activation. We also confirmed this by flow cytometry analysis of immune cell populations in GWAT (Fig. 5B), where we found more macrophages and T cells in the KO animals compared to controls, especially more pro-inflammatory polarized F4/80+ CD11c+ macrophages (Fig. 5C,D). In line, with the *in vitro* data, this was also linked to higher gene expression of ER stress surrogate marker genes such as *Atf3*, *Atf4, Atf6* and *Ddit3* as well as inflammatory marker *Ccl2* and *Cd68* (Fig. 5E).

**Fig. 5:**
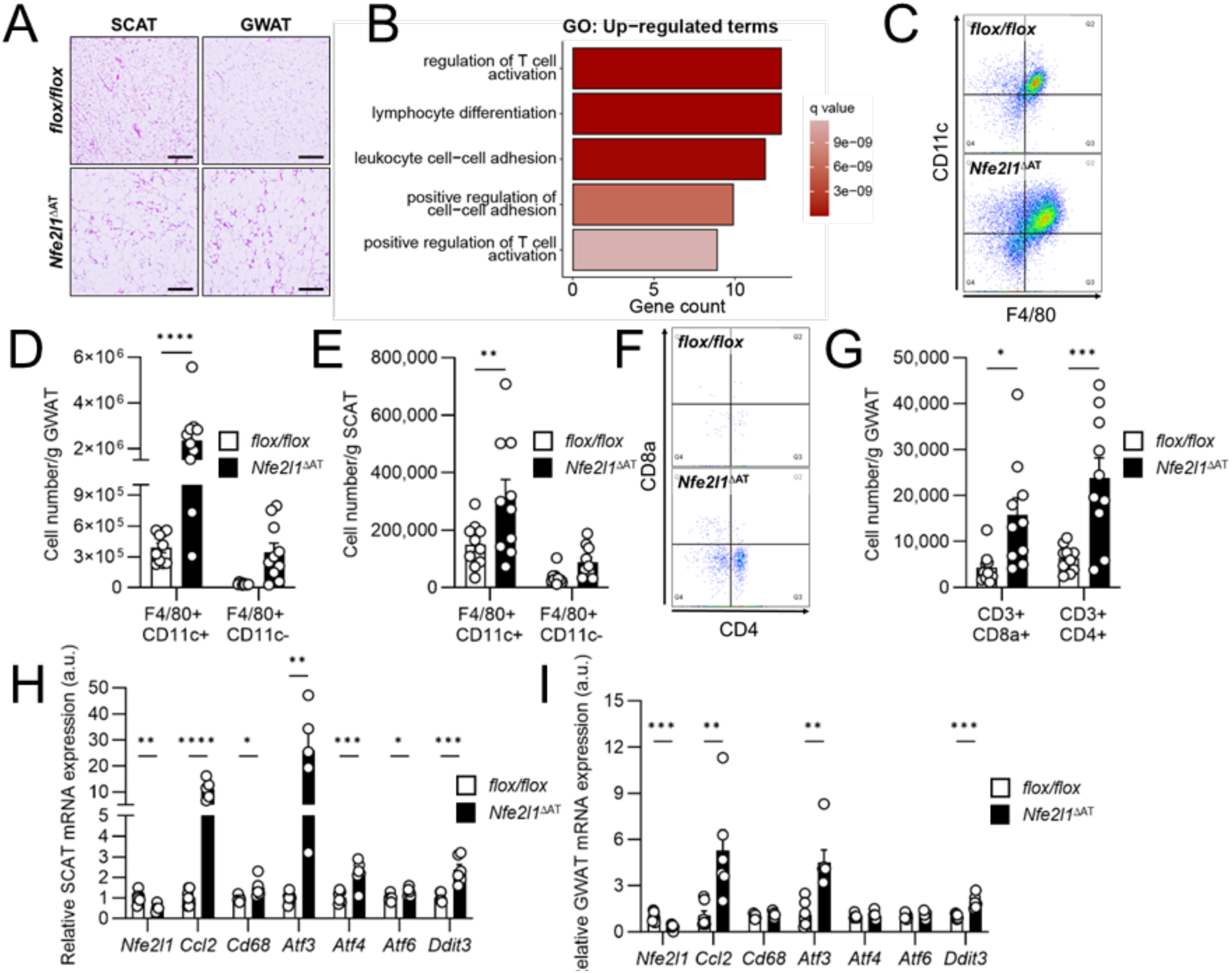
Loss of adipocyte Nfe2l1 induces inflammation in WAT. (A) Representative hematoxilin-eosin staining of SCAT and GWAT from *flox/flox* and *Nfe2l1*^ΔAT^ mice (scale bar: 200 µm). (B) Up-regulated GO terms in *Nfe2l1*^ΔAT^ SCAT vs. *flox/flox* SCAT, n=4. (C) Representative flow cytometry blot of adipose tissue stellate vascular fraction (SVF) stained for F4/80 and CD11c. (D-E) Absolute cell number of F4/80+ cells in GWAT and in SCAT from *flox/flox* and *Nfe2l1*^ΔAT^ mice, n=10, data are mean+s.e.m, \*\**P*<0.01, \*\*\*\**P*<0.001 tested by 2-way ANOVA with Šidák post-hoc test. (F) Representative flow cytometry blot of adipose tissue SVF stained for CD4 and CD8a. (G) Absolute cell number of CD3+ cells in GWAT from *flox/flox* and *Nfe2l1*^ΔAT^ mice, n=10, data are mean+s.e.m, \**P*<0.05, \*\*\**P*<0.001 tested by 2-way ANOVA with Šidák post-hoc test. (H-I) relative mRNA levels of inflammation and stress associated genes in SCAT and GWAT from *flox/flox* and *Nfe2l1*^ΔAT^ mice, n=5-7, data are mean+s.e.m, \**P*<0.05, \*\**P*<0.01, \*\*\**P*<0.001, \*\*\*\**P*<0.0001 tested by 2-way ANOVA with Šidák post-hoc test.

As inflammation of the adipose tissue is a hallmark of insulin resistance and atherosclerosis, we investigated the impact of Nfe2l1 in white adipocytes for obesity and associated metabolic dysfunction. For that, we fed *Nfe2l1*^ΔAT^ and *flox/flox* mice a 60 % fat HFD for 16 weeks. We did not observe differences in body weight or baseline energy expenditure between *flox/flox* and *Nfe2l1*^ΔAT^ mice (Supplementary Fig. 2). However, *Nfe2l1*^ΔAT^ mice on DIO exhibited a re-distribution of body mass from white fat mass to liver mass (Fig. 6B,C). We found that on HFD, *Nfe2l1*^ΔAT^ were less glucose-tolerant and more insulin-resistant than *flox/flox* mice (Fig. 6D,E). Moreover, in line with lower adipose mass, *Nfe2l1*^ΔAT^ mice had lower leptin (Fig. 6F) and adiponectin plasma levels (Fig. 6G) but higher insulin (Fig. 6H). Notably, we also observed lower adiponectin plasma level in the absence of any differences in body or tissue weights in *Nfe2l1*^ΔAT^ mice on chow diet (Tab. 1). In summary, in the absence of Nfe2l1, adipocyte health was compromised and resulted in insulin resistance and metabolic dysfunction. Although transient loss of Nfe2l1 had only a minor effect on cholesterol homeostasis in cultured adipocytes, we still investigated whether in mice loss of adipocyte Nfe2l1 is linked to cholesterol-induced cardiometabolic diseases using the Apolipoprotein E (ApoE)-deficient mouse. These mice are very sensitive to dietary cholesterol and develop hypercholesterolemia and complex inflammation on a Western-type diet (WD)^28^.

**Tab. 1:** Overview of plasma parameter. Data are mean±s.e.m., Comparison of genotypes on the same diet, \**P*<0.05, \*\**P*<0.01, \*\*\**P*<0.001, \*\*\*\**P*<0.0001 by Student‘s T-test and Mann-Whitney U-test, where appropriate.

| Plasma parameter | Triglycerides (mg/dL) | Cholesterol (mg/dL) | Insulin (ng/mL) | Adiponectin (μg/mL) | Leptin (ng/mL) |
| --- | --- | --- | --- | --- | --- |
| *flox/flox* chow (n=6) | 60 ± 6 | 125 ± 2 | 0.2 ± 0.0 | 9.2 ± 0.6 | 14.4 ± 3.9 |
| *Nfe2l1*<sup>ΔAT</sup> chow (n=6) | 51 ± 3 | 124 ± 7 | 0.3 ± 0.0* | 4.3 ± 0.3**** | 11.2 ± 5.6 |
| *flox/flox* DIO (n=7) | 43 ± 3 | 172 ± 7 | 1.17 ± 0.03 | 9.9 ± 0.1 | 130.9 ± 5.5 |
| *Nfe2l1*<sup>ΔAT</sup> DIO (n=8) | 45 ± 5 | 160 ± 5 | 1.48 ± 0.06** | 7.0 ± 0.2* | 94.0 ± 5.8* |
| *Apoe*<sup>-/-</sup>*flox/flox* chow (n=8) | 47 ± 12 | 508 ± 37 | 0.9 ± 0.2 | 4.7 ± 0.3 | 3.4 ± 0.4 |
| *Apoe*<sup>-/-</sup>*Nfe2l1*<sup>ΔAT</sup> chow (n=8) | 116 ± 11* | 528 ± 30 | 0.9 ± 0.2 | 2.2 ± 0.1* | 3.0 ± 0.4 |
| *Apoe*<sup>-/-</sup>*flox/flox* WD (n=9) | 105 ± 12 | 760 ± 78 | 0.3 ± 0.0 | 10.8 ± 0.5 | 12.6 ± 2.7 |
| *Apoe*<sup>-/-</sup>*Nfe2l1*<sup>ΔAT</sup> WD (n=12) | 126 ± 11 | 996 ± 23** | 1.4 ± 0.4** | 2.2 ± 0.2**** | 4.5 ± 0.4*** |

**Fig. 6:**
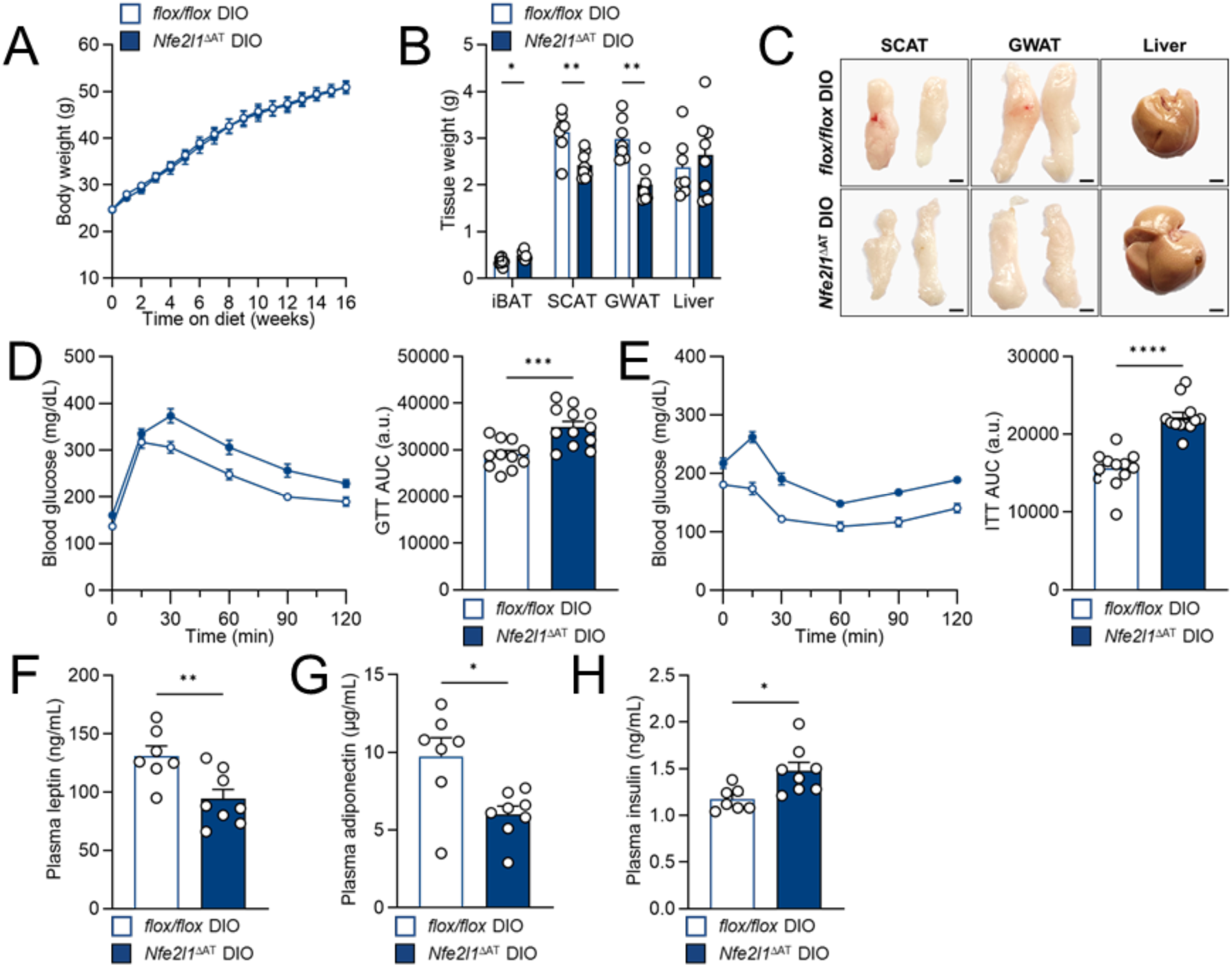
Adipocyte Nfe2l1 confers tolerance to diet-induced obesity. (A) Body weight of *flox/flox* and *Nfe2l1*^ΔAT^ mice on high-fat diet (HFD), n=14, data are mean+s.e.m. (B) Tissue weights, n=8, data are mean + s.e.m., \**P*<0.05, \*\**P*<0.01 by Student’s T-test. (C) Representative pictures of tissue samples *ex vivo*. (D) i.p. GTT and area under the curve (AUC), n=12, data are mean+s.e.m., \*\*\**P*<0.001 by Student’s T-test. (E) i.p. ITT and AUC, n=12, data are mean + s.e.m., \*\*\*\**P*<0.0001 by Student’s T-test. (F-H) Plasma insulin, adiponectin, and leptin levels, n=6-8, data are mean + s.e.m., * by Student’s T-test.

Thus, we crossed *Nfe2l1*^ΔAT^ onto an ApoE-deficient background (*Apoe*^−/−^ *Nfe2l1*^ΔAT^) and fed these mice a WD to induce atherosclerosis. Somewhat in line with the DIO model, we observed a lipoatrophy-like phenotype in the absence of any body weight differences (Fig. 7A), characterized by markedly lower adipose tissue mass, especially in the SCAT depot (Fig. 7B,C). Compared to their *Apoe*^−/−^ *flox/flox* littermates, *Apoe*^−/−^ *Nfe2l1*^ΔAT^ mice were equally glucose-tolerant (Fig. 7D) but more insulin-resistant (Fig. 7E), and had higher plasma levels of cholesterol and insulin, as well as lower adiponectin and leptin plasma levels (Tab. 1). This was linked to enhanced atherogenesis, dyslipidemia, and systemic inflammation in *Apoe*^−/−^*Nfe2l1*^ΔAT^ mice compared to *Apoe*^−/−^ *flox/flox* littermates on WD (Fig. 7F-I). Interestingly, the lipoatrophy was specific for the loss of Nfe2l1 in white adipocytes, as we observed lower WAT mass in *Nfe2l1*^ΔAT^ mice but not in uncoupling protein 1 (Ucp1)-Cre Nfe2l1 KO mice on WD (referred to as *Nfe2l1*^ΔBAT^, Supplementary Fig. 3). In summary, these *in vivo* studies show that Nfe2l1 is linked to adipose inflammation, insulin resistance, dyslipidemia, and atherosclerosis.

**Fig. 7:**
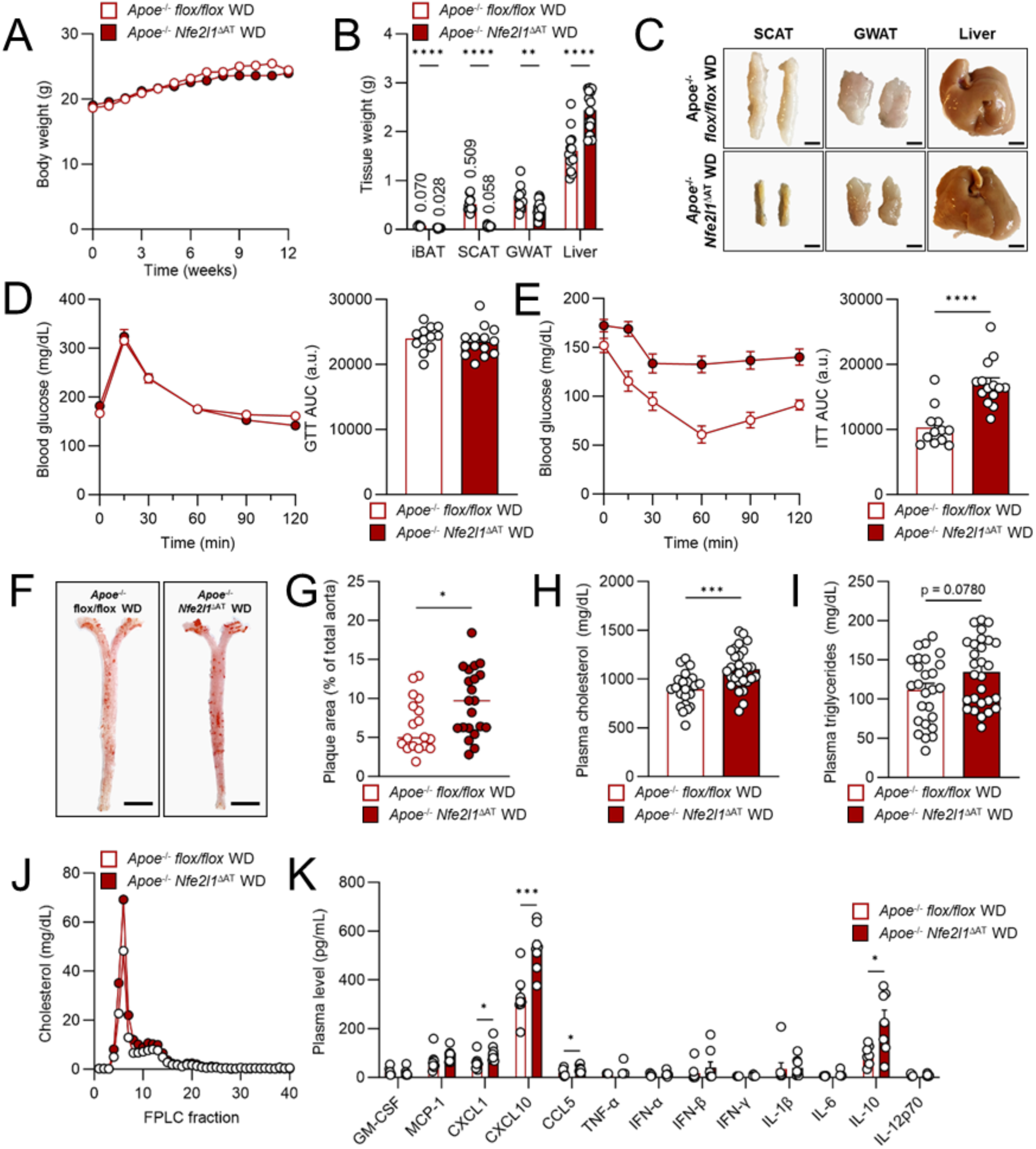
Adipocyte Nfe2l1 protects from systemic inflammation, dyslipidemia and atherosclerosis. (A) Body weight of *Apoe*^−/−^ *flox/flox* and *Apoe*^−/−^ *Nfe2l1*^ΔAT^ mice on Western-type diet (WD), n=14-17, data are mean + s.e.m. (B) Tissue weights, n=13-14, data are mean+s.e.m., \*\**P*<0.01, \*\*\*\**P*< 0.001 by Student‘s T-test. (C) Representative tissue pictures *ex vivo*. (D, E) i.p. GTT, i.p. ITT and their and AUC of *Apoe*^−/−^ *flox/flox* and *Apoe*^−/−^ *Nfe2l1*^ΔAT^ mice on WD, n=12-14, data are mean+s.e.m., \*\*\*\**P*<0.0001 by Student’s T-test. (D) Representative pictures of *en face* stained aortas (scale bar: 0.5 cm). (E) Quantification of plaque area, n=18-21, data are mean+s.e.m., \**P*<0.05 by Student‘s T-test. (F) Plasma cholesterol, n=24-29, data are mean + s.e.m., \*\*\**P*<0.001 by Student‘s T-test. (G) Plasma triglycerides, n=24-29, data are mean+s.e.m., *P* by Mann-Whitney U test. (H) FPLC of pooled plasma, each generated by pooling of n=8 individual samples. (I) Cytokine and chemokine plasma concentrations, n=7-8, data are mean+s.e.m., \**P*<0.05, \*\*\**P*< 0.001 by Student‘s T-test.

## Discussion

The adaptation of adipocytes to excess nutrients, particularly lipids, is a pivotal process for maintaining systemic metabolic homeostasis. Failure to facilitate adequate adipose tissue expansion leads to adipocyte stress, tissue inflammation, and ectopic lipotoxicity. This is the turning point when adipocytes become maladaptive and contribute to the progression of cardiometabolic diseases. The ER plays a critical role for the metabolic adaptation of cells by nutrient sensing and regulating adaptive stress resistance pathways. The activation of stress-resistance pathways, notably the UPR, is well documented in obese, insulin-resistant tissues, especially adipose tissue, where ER stress contributes to adipocyte dysfunction^11,29,30^. However, little is known about the mechanisms that protect adipocyte health in obesity. Here, we show that the Nfe2l1-proteasome pathway is a critical pillar of adipocyte health. In this way, Nfe2l1 mediates the adaptation to lipid-rich diets and protects adipocytes from lipotoxicity, particularly cholesterol-induced inflammation, which resulted in insulin resistance and atherosclerosis in Nfe2l1-deficient animals.

Nfe2l1 adaptively activates the transcription of proteasome subunit genes when proteasomal activity is compromised^23,24,31^. Such precise fine-tuning in proteasome regulation is vital for survival, highlighted by the lethality of global Nfe2l1 deletion during embryonic development^32^. Tissue-specific deletion of Nfe2l1 underscores its necessity for maintaining normal tissue function by regulating metabolism and inflammation under metabolic stress conditions in BAT^16^, liver^12–14^, heart^15^ and now we extend this paradigm to the role of Nfe2l1 in white adipocytes. The phenotypic outcomes of Nfe2l1 deficiency should be interpreted as tissue-specific consequences of compromised proteasome function. At the cellular level, white adipocytes lacking Nfe2l1 showed the expected impairment of proteasome activity, which was linked to the activation of pro-inflammatory stress kinase signaling pathways. In humans, monogenetic diseases caused by mutations in proteasome subunits are referred to as proteasome-associated auto-inflammatory syndromes, which have a strong inflammatory component, and we have recently shown that manipulation of these *PSM* genes causes inflammation in adipocytes^20^. A major similarity between our previous studies and this current work is that direct inhibition of the proteasome by chemical inhibitors or silencing of structural components provokes a much stronger phenotype *in vitro* than does silencing of Nfe2l1. This emphasizes the adaptive nature of Nfe2l1 and only its chronic absence, e.g. as observed in the mouse models here and previously^16^, results in a progressive phenotype, which is strongly enhanced by lipid-rich dietary challenges.

As we observed a more pronounced lipodystrophy-like phenotype in mice on cholesterol-enriched WD, we further investigated the response of adipocyte proteostasis to cellular cholesterol loading. Silencing of Nfe2l1 in adipocytes by itself had little impact on inflammation or stress marker genes but cholesterol treatment induced stress and inflammation markers, which was synergistically escalated by the combination of both cholesterol and chemical proteasome inhibition by epoxomicin. Cholesterol treatment significantly altered the adipocyte transcriptome towards a lipotoxic stress response characterized by inflammation, ER stress, and disrupted lipid metabolism. Atf3 was the most highly induced transcription factor by cholesterol treatment and was also highly induced in adipose tissue when Nfe2l1 was deleted. As the cholesterol-induced lipotoxic response was significantly attenuated by silencing of Atf3 it stands to reason that the activation of Atf3 induces transient inflammation but remains elevated in the chronic absence of Nfe2l1. In addition to that, we also investigated whether cholesterol activated stress kinase and inflammatory signaling pathways, a feature observed in Nfe2l1 KO adipocytes, and if silencing of Atf3 would be able to diminish or even block it. Interestingly, while cholesterol treatment activated NF-κB, JNK and p38 signaling, which was even exacerbated by the addition of epoxomicin, silencing of Atf3 did not diminish JNK and NF-κB signaling in 3T3-L1 adipocytes. Only p38 activation in response to cholesterol treatment was blunted by silencing of Atf3, which suggests p38 as an additional player between Atf3 and inflammatory gene transcription. Also, this means that adipocytes possess networks of distinct cellular inflammation pathways.

Obesity and atherosclerosis are immunometabolic diseases, which are linked the activation of pro-inflammatory signaling pathways and the irregular production of cytokines and chemokines^3,10,33^. We found that the absence of Nfe2l1 exacerbates inflammation in WAT, evidenced by the marked infiltration of F4/80+ CD11c+ macrophages and CD4+ and CD8+ T cells. While CD4+ T cells typically inhibit macrophage infiltration by secreting Il-4 and Il-10, CD8+ T cells initiate and drive inflammation in obese adipose tissue by promoting macrophage activation and migration^34,35^. Inflammation is a driver of atherosclerosis and promotes its medical complications^28^. Inflammation in perivascular AT is considered a risk factor for the development of atherosclerosis, as it is located in close proximity to the vessel wall and has paracrine effects there^36–38^. Although secretion of pro-inflammatory mediators such as TNFα and MCP-1 from adipocytes is well established, the contribution of adipocytes especially to circulating pro-inflammatory cytokines and chemokines remains unclear. In this study, loss of adipocyte Nfe2l1 was directly linked to enhanced atherogenesis, potentially caused by dyslipidemia and higher level of circulating pro-inflammatory cytokines Cxcl1, Ccl5 and Cxcl10. NF-κB-mediated secretion of Cxcl10 from adipocytes and infiltration of T cells into AT^39^ might thus explain the substantial occurrence of CD8+ and CD4+ T cells in Nfe2l1-deficient WAT. *In vitro*, stimulation of adipocytes with cholesterol induced the expression of *Cxcl1* and *Cxcl10*, indicating that disturbed proteasomal proteostasis causally contributes to circulating cytokine levels.

To investigate the impact of adipocyte Nfe2l1 in the context of obesity and cardiometabolic disorders, we established *Nfe2l1*^ΔAT^ mice on both a standard C57BL6/J and an *Apoe*^−/−^ genetic background and then challenged them with lipid-rich diets. While DIO and an atherogenic WD resulted in distinct phenotypes, both diets led to lower SCAT and GWAT mass. Notably, WD also led to lower BAT mass, which was not observed with a standard chow diet or in DIO. SCAT is the primary lipid storage in the body and only when its storage capacities exceed, lipids accumulate in the VISAT^40^. Thus, lower SCAT mass indicates a reduced ability to safely store lipids, which leads to ectopic lipid deposition in the presence of excessive nutrient intake^6^. A consistent outcome of reduced WAT mass was the lower plasma levels of adiponectin and leptin in the absence of Nfe2l1, the predominant hormones produced by AT, which altogether indicates compromised adipocyte function and health. This lipodystrophy-like phenotype also resulted in elevated hepatic lipid deposition and in *Apoe*^−/−^ *Nfe2l1*^ΔAT^ mice enhanced dyslipidemia, which is especially important in the context of atherosclerosis, in which circulating atherogenic lipoproteins initiate the disease^28,41^. The elevated hepatic and plasma lipids are presumably consequences of insufficient metabolic buffering of dietary cholesterol by the adipocytes, as WAT mass was severely diminished on WD. Finally, lower circulating adiponectin level *per se* might also enhance atherosclerosis, since adiponectin is anti-inflammatory and anti-fibrotic activities^42^ and its overexpression is sufficient to lower atherogenesis^43^.

In a human cohort over a broad range of BMI, lower *NFE2L1* WAT expression was associated with higher BMI, and almost all proteasome subunit genes were positively correlated to *NFE2L1* gene expression. This indicates that the entire Nfe2l1-proteasome pathway is downregulated in obese humans. Thus, it is tempting to speculate that reactivating or enhancing the Nfe2l1-proteasome pathway could help to improve adipocyte dysfunction and insulin resistance. However, to our experience this is a challenging task: Ways to enhance proteasome function by small molecules are limited and while genetic approaches work in cells^44,45^, they may lack clinical efficacy. In conclusion, our work highlights the complex interplay of genetic predisposition and dietary lipids that dictates the diverse outcomes of cardiometabolic disease. We show that the Nfe2l1-proteasome pathway is essential for white adipocytes to adjust to lipid-rich diets, safeguarding adipocytes against lipotoxicity, inflammation, and systemic metabolic disorders.

## Methods

### Human data

The human data utilized in this study were obtained from the Leipzig Obesity Biobank (LOBB), which includes paired samples of abdominal SCAT and omental VISAT, as well as body fluids and associated anthropometric data. During elective laparoscopic abdominal surgeries, adipose tissue samples were acquired following established protocols^46^. Body composition and metabolic parameters were evaluated using standardized methods as outlined in previous studies^47,48^. The study received approval from the Ethics Committee of the University of Leipzig (Approval no: 159-12-21052012) and was conducted in compliance with the Declaration of Helsinki principles. All participants provided written informed consent prior to their inclusion in the study. Exclusion criteria encompassed participants under 18 years of age, chronic substance, or alcohol misuse, smoking within the 12 months prior to surgery, acute inflammatory diseases, use of glitazones as concomitant medication, end-stage malignant diseases, weight loss exceeding 3 % in the three months preceding surgery, uncontrolled thyroid disorder, and Cushing’s disease. The first cohort consisted of 478 individuals, categorized into non-obese (n=68; age: 62.7±17.4 years old; BMI: 24.4±2.9 kg/m^2^) and obese (n=410; age: 46.6±11.4 years old; BMI: 49.7±9.0 kg/m^2^) groups. *NFE2L1* expression in SCAT and VISAT was quantified using quantitative real-time polymerase chain reaction (qRT-PCR), and the expression levels were normalized to *HPRT1*. The second cohort comprised 31 non-obese individuals (Age: 55.8±13.4 years old; BMI: 25.7±2.7 kg/m^2^) and 1,448 obese individuals (Age: 46.9±11.7 years old; BMI: 49.2±8.3 kg/m^2^). In this cohort, gene expression analysis of *NFE2L1* and proteasome subunit genes was performed by RNA-Seq.

### Animal care and use

All animal experiments were performed according to procedures approved by the animal welfare committees of the government of Upper Bavaria, Germany (ROB-55.2-2532.Vet_02-20-32) and performed in compliance with German Animal Welfare Laws. We housed animals in individually ventilated cages at room temperature (22 °C) with a 12 h light–dark cycle and *ad libitum* access to standard chow diet (Ssniff) and water. The previously described Nfe2l1 floxed mice^16^ were crossed with B6.FVB-Tg(Adipoq-cre)1Evdr/J mice (The Jackson Laboratory, stock no. 028020) to generate Nfe2l1 adipocyte-specific knockout (*Nfe2l1*^ΔAT^) mice. We used Cre negative (Cre^wt/wt^) littermates carrying floxed Nfe2l1 alleles as controls (*flox/flox*). For atherosclerosis studies we have crossed Adipoq-Cre Nfe2l1 mice with whole-body Apoe knock-out mice (*Apoe^−/−^*, The Jackson Laboratory, stock no. 002052), to create *Apoe^−/−^ Nfe2l1*^ΔAT^ and *Apoe^−/−^ flox/flox* controls. We also studied brown-adipocyte specific Nfe2l1 ablation using Ucp1-Cre Nfe2l1 KO mice (*Nfe2l1*^ΔBAT^), as previously described^16^. For DIO studies, we fed male mice High-fat diet 60 kJ% fat (E15742-34, Ssniff) for 16 weeks starting at age 4-8 weeks. For atherosclerosis studies, we fed female mice Western diet (E15721-34, Ssniff) containing 42 kJ% fat enriched with 0.21 % cholesterol for 12 weeks. We measured respiration, energy expenditure, food and water intake, as well as activity using a Promethion Core system (Sable Systems). BAT-specific energy expenditure was measured by subcutaneous injection of CL316,243 (CL, Tocris, 0.5 mg/kg in 0.9 % w/v in NaCl) at ∼12:00 p.m. We assessed glucose and insulin tolerance with i.p. GTT and i.p. ITT. For both GTT and ITT, mice were starved for 6 h, baseline blood glucose was measured and then mice were injected i.p. with glucose (1 g/kg body weight, Gibco) and insulin (0.25-0.75 U/kg body weight, Abbott, in DPBS + 0.2 % BSA), respectively. Blood glucose was subsequently measured at 15 min, 30 min, 60 min, 90 min and 120 min after injection. At the end of the study, mice were anesthetized with a lethal dose of ketamine-xylazin (120 mg/kg and 8 mg/kg body weight, respectively). After ensuring deep anesthesia, the chest of the mouse was disinfected, opened, and blood was collected via cardiac puncture. We perfused with cold DPBS with 50 U/mL heparin, dissected the organs and weighed them. Organs were snap frozen in liquid nitrogen and stored at −80 °C. We estimated sample size by pilot experiments that showed trends of effects and their sizes. In most cases, n=6 was the minimum number of mice used. During feeding studies, we excluded mice for poor body condition or barbering. All experiments were reproduced at least in two independent cohorts. Genotyping of the Nfe2l1 floxed mouse model and recombination of the *Nfe2l1* gene was assessed as described previously^16^.

### Aorta dissections and en face Oil Red O staining

After cardiac blood collection and perfusion with cold DPBS, gastrointestinal tract, liver, lung and thymus were removed to expose the aorta. Perivascular adipose tissue was carefully removed *in situ* under a stereomicroscope using Vannas spring scissors and Dumont forceps. We harvest the aorta, including parts of the brachiocephalic and subclavian arteries, and transferred it on a black pinning bed. The remaining adipose tissue was removed, the aorta was split longitudinally and pinned down with minutiae needles. Pinned aortas were fixed in zinc formalin (Sigma) overnight and transferred to DBPS (Gibco) the next day. For the staining, we diluted Oil-Red-O stock solution (Sigma) to 60 % with ddH2O (v/v) and filtered it through a 0.22 µm syringe filter. We first dipped the pinned aorta in 60 % 2-propanol (v/v, 10x) and then stained it with 60 % Oil Red O staining solution (v/v) for 15 min. Excessive stain was removed by dipping in 60 % 2-propanol (v/v, 10x) and aortas were transferred to ddH2O until imaging. We took macroscopic pictures using a Canon EOS 4000D. Total aorta area and plaque area were quantified using ImageJ. For long-term conservation, aortas were mounted on microscopy slides with Kaiser’s glycerol (Roth).

### Histology

Tissues were collected in histology cassettes, fixed in zinc formalin overnight (12-24 h) and dehydrated by successively immersing the tissues in 70 % ethanol (v/v), 96 % ethanol (v/v), 100 % ethanol (v/v), and Xylene (Sigma) in an ASP200S (Leica) apparatus The tissues were embedded in paraffin (Roth) sliced in 4-5 μm sections using a microtome (Leica). The sections underwent deparaffination and rehydration procedures, by dipping the slides in 2x Xylene (5 min), 2× 100 % ethanol (v/v, 5x), 96 % ethanol (v/v, 5x), 70 % ethanol (v/v, 3x), and ddH_2_O (10x). We stained sections with hematoxylin (5 min), washed with ddH_2_O and then stained with eosin (5 min), followed by dehydration with 3x 70 % ethanol (v/v), 5x 96 % ethanol (v/v), 5× 100 % ethanol (v/v), 5× 100 % ethanol (v/v) and 2x Xylene (5 min). The slides were mounted with Histokitt II (Roth), images taken with a DMi8 (Leica).

### Plasma analysis

Whole blood was collected by heart puncture and transferred to EDTA blood collection tubes. Plasma was prepared by centrifuging at 2,000 *g* for 10 min at 4 °C. We used Mouse/Rat Leptin Quantikine ELISA Kit (R&D systems), Mouse Adiponectin/Acrp30 Quantikine ELISA Kit (R&D systems) and Ultra-Sensitive Mouse Insulin ELISA Kit (Crystal Chem) according to manufacturer’s instructions to quantify plasma leptin, adiponectin and insulin level, respectively. Plasma cholesterol and triglyceride level were measured using Cholesterol FS and Triglyceride FS kits (Diasys) according to manufacturer’s instructions. We assessed pro- and anti-inflammatory cytokine and chemokine concentrations using the LEGENDplex™ Mouse Anti-Virus Response Panel (Biolegend), which was analyzed on a MACSQuant® Analyzer 10 (Miltenyi).

### Flow cytometry analysis of white adipose tissue SVF

For flow cytometry analysis of immune cell population in WAT, we dissected GWAT and minced it with surgical scissors. To each fat pad we added DMEM/F-12 (Sigma-Aldrich), 1 % PenStrep (Sigma-Aldrich), 15 mg/mL fatty acid free BSA (Sigma-Aldrich), 1 mg/ml collagenase type 2 (Worthington) and 0.1 mg/mL DNase 1 (Roche) and incubated at 37 °C in a thermoshaker for approx. 30 min, shaking at 800 rpm with manually shaking every 10 min. The digestion was stopped by adding staining buffer (DPBS, 1 % FBS v/v, 2 mM EDTA, 0.1 % sodium azide w/v) in a 1:1 ratio. SVF were filtered through a 70 µm filter and centrifuged at 500 x g for 5 min (4 °C). Red blood cells (RBC) were lysed by incubation with RBC lysis buffer (Qiagen) for 3 min at room temperature. We stopped the lysis by adding staining buffer in a 5:1 ratio and centrifuged at 500 *g* for 5 min (4 °C). The cell pellet was re-suspended in staining buffer supplemented with 1:100 anti-mouse CD16/32 (Fc block, BioLegend) and incubated on ice for 10 min. After centrifugation at 500 *g* for 5 min (4 °C), supernatant was decanted. Cells were stained for 30 min in the dark (4 °C) with either anti-CD45.2, anti-CD11b, anti-F4/80 and anti-CD11c fluorescence-labeled antibodies for macrophages or anti-CD45.2, anti-CD11b, anti-CD19, anti-Cd8a, anti-CD4 fluorescence-labeled antibodies for T cells (Supplementary Tab. 1). Cells were washed by adding staining buffer in a 4:1 with subsequent centrifugation 500 *g* for 5 min (4 °C). After removing the supernatant, we re-suspended the pellet in staining buffer, added countbright beads (Invitrogen) and mixed by vortexing for 30 s. After centrifugation at 500 *g* for 5 min (4 °C), pellets were re-suspended in staining buffer and analyzed on a MACSQuant® Analyzer 10 (Miltenyi). Data was analyzed using FlowJo (v.10). We corrected for spectral overlap by applying compensation values derived from single stained samples and assured correcting gating with Fluorescence Minus One controls. The gating strategies can be found in Supplementary Figure 4. We calculated absolute cell numbers according to countbright beads (Invitrogen) instructions and normalized cell number to tissue weight.

### Cell culture and treatments

We used 3T3-L1 cells (Sigma), which were differentiated into mature white adipocytes. The cells were induced at confluence for 2 days and differentiated for 2 more days followed by 1 day of cultivation in standard medium (DMEM GlutaMax (Gibco), 10 % iron-fortified calf serum (Sigma-Aldrich), 1 % penicillin-streptomycin (Sigma-Aldrich)). For both induction and differentiation, the standard medium was supplemented with 850 nM human insulin (Sigma-Aldrich), 1 µM dexamethasone (Sigma-Aldrich, in 100 % ethanol), 1 µM rosiglitazone (Cayman Chemicals, in 100 % DMSO) and 500 µM IBMX (Sigma-Aldrich, in 100 % DMSO). All treatments were performed on the fifth day of differentiation. Proteasome inhibitor epoxomicin (Millipore) was used at 100 nM in 100 % DMSO. For cholesterol treatments, cholesterol powder (Sigma) was dissolved in 100 % (v/v) ethanol at 65 °C and conjugated to methyl-beta-cyclodextrin (MbCD) at a concentration of 5 mM cholesterol to 42 mg/ml MbCD. The conjugation to MbCD keeps cholesterol solubilized and causes the cells to accumulate cholesterol, as indicated by down-regulation of Srebp2 target genes^12^.

### Primary cell preparation and culture

For primary cell experiments, mature adipocytes were differentiated from pre-adipocytes, isolated from the SVF of adipose tissue of 4 weeks old C57BL/6J (Janvier) wild-type mice. The mice were sacrificed by cervical dislocation, the respective adipose tissues were harvested and pooled. The tissues were minced, weighted and digested in DMEM/F-12 (Sigma-Aldrich), 1 % PenStrep (Sigma-Aldrich), 15 mg/mL fatty acid free BSA (Sigma-Aldrich), 1 mg/mL collagenase type 2 (Worthington) and 0.1 mg/ml DNase 1 (Roche) for 30-45 min at 37 °C. For BAT, the digestion mix was supplemented with 1.2 U/ml Dispase (Roche). The digestion was stopped by adding DMEM/F-12 (+10 % FBS, +1 % PenStrep) in a 1:5 ratio. The digest was filtered through a 100 µm strainer and centrifuged at room temperature for 10 min at 500 *g*. The mature adipocytes and the supernatant were aspirated, the pellet suspended in DMEM/F-12 (+10 % FBS, +1 % PenStrep) and filtered through a 70 µm and a 30 µm strainer before plating in a T75 flasks. The medium was changed the next day, and then every other day until the cells reached confluency. To differentiate pre-adipocytes into mature adipocytes, cells were induced for 2 days and differentiated for 2 days followed by 1 day cultivation in standard medium. The induction medium consisted of DMEM/F-12 (+10 % FBS, +1 % PenStrep) supplemented with 340 nM human insulin (Sigma-Aldrich), 1 µM dexamethasone (Sigma-Aldrich, in 100 % ethanol), 1 µM T3 (Sigma-Aldrich, in 1 M NaOH), 1 µM rosiglitazone (Cayman Chemicals, in 100 % DMSO), 500 µM IBMX (Sigma-Aldrich, in 100 % DMSO). The differentiation medium for white adipocytes from the SCAT consisted of DMEM/F-12 (+10 % FBS, +1 % PenStrep) with 10 nM human insulin and 2 µM T3.

### Reverse transfection and RNAi

To knock-down genes of interest, we reverse transfected 3T3-L1 and primary white adipocytes with SMARTpool siRNA (Dharmacon) and Lipofectamine™ RNAiMAX transfection reagent (Thermo Fisher Scientific) on the third day of differentiation. In brief, we diluted RNAiMAX and SMARTpool siRNAs, respectively, with Opti-MEM® (Gibco) according to RNAiMAX manufacturer’s instructions and incubated for 5 min at room temperature. SMARTpool siRNAs for *Nfe2l1* and *Atf3* were used in a final concentration of 30 nM for single knockdowns. For double knockdown of *Nfe2l1* and *Atf3*, SMARTpool siRNAs were mixed in an equimolar ratio with final concentrations of 60 nM. Afterwards, diluted Lipofectamine® RNAiMAX was mixed 1:1 with diluted SMARTpool siRNAs and incubated for another 20 min. In the meantime, cells were trypsinized, centrifuged and the pellet re-suspended in the appropriate amount of differentiation medium. Finally, the siRNA-RNAiMAX mix was distributed in a cell culture plate and the adipocytes were pipetted on top. Careful shaking of the plate ensured an even distribution of the reaction mixture and the cells were allowed to reattach at room temperature for approximately 30 min before being transferred to the incubator. After 24 h, the transfection mix was replaced with the differentiation medium and the cells were incubated for another 24 h, after which we treated or directly harvested them.

### RNA extraction, cDNA synthesis and qPCR

To extract total RNA from frozen adipose tissue and cells, we used the NucleoSpin® RNA kit (Macherey-Nagel). Cells were lysed by adding RA1 supplemented with 1 % β-mercaptoethanol (Sigma), whereas tissue samples were lysed with TRIzol reagent (Thermo Fisher Scientific) in a TissueLyser II at 30 hz for 3 min. Cell lysates were filtered using a filter column (Macherey-Nagel) and the flow-through was mixed with 70 % ethanol (v/v) in a 1:1 ratio. The tissue lysates were supplemented with 100 % chloroform (Sigma) in a 1:5 ratio (v/v), mixed and centrifuged for 15 min at 4 °C and maximum speed. The resulting clear phase was collected and also mixed with 70 % ethanol (v/v) in a 1:1 ratio. From here on, the lysate-ethanol mixture was applied to RNA columns (Macherey-Nagel) and the RNA was further processed according to the manufacturer’s instructions. RNA concentrations were measured on a NanoDrop spectrophotometer (Thermo Fisher Scientific). To synthesize complementary DNA (cDNA), we reverse-transcribed 500 ng RNA with the Maxima™ H Master Mix 5x (Thermo Fisher Scientific) in a total volume of 10 μl according to manufacturer’s instructions (10 min 25 °C, 15 min 50 °C, 5 min 85 °C). cDNA was diluted 1:40 with RNase-free H_2_O. Relative gene expression was quantified using quantitative real time-polymerase chain reaction (qPCR). Each reaction contained 4 µL cDNA, 5 µL PowerUp™ SYBR Green Master Mix (Applied Biosystems) and 1 µL of 5 µM primer stock (Supplementary Tab. 2). We used standard run conditions for Applied Biosystems SYBR Green Gene Expression Assays (2 min 50 °C, 10 min 95 °C, 40 cycles of 0:15 min 95 °C, 1 min 60 °C). Cycle thresholds of genes of interest were normalized to *Tbp* levels by the ΔΔCt-method and displayed as relative copies per *Tbp* or relative expression normalized to experimental control groups.

### Protein extraction and Western blotting

We lysed cells in RIPA buffer (150 mM NaCl (Merck), 5 mM EDTA (Merck), 50 mM Tris pH 8 (Merck), 0.1 % w/v SDS (Carl Roth), 1 % w/v IGEPAL® CA-630 (Sigma-Aldrich), 0.5 % w/v sodium deoxycholate (Sigma-Aldrich)) freshly supplemented with complete EDTA protease inhibitors (Roche) and PhosStop (Roche). The lysates were centrifuged at 4 °C for 30 min at 21,000 *g* to clear from debris and lipids. Protein concentrations were determined using the Pierce BCA Protein Assay (Thermo Fisher Scientific) according to the manufacturer’s instructions. For western blotting, lysates were adjusted to a final concentration of 20-30 µg/lane in 1x Bolt™ LDS Sample buffer (Thermo Fisher Scientific) supplemented with 5 % (v/v) 2-mercaptoethanol (Sigma-Aldrich). For SDS-PAGE, we used Bolt™ 4-12 % Bis-Tris gels (Thermo Fisher Scientific) with Bolt™ MOPS SDS running buffer. After separation, proteins were transferred onto a 0.2 µm PVDF membrane (Bio-Rad) using the Trans-Blot® Turbo™ system (Bio-Rad) at 12 V, 1.4 A for 16 min. The membranes were briefly stained with Ponceau S (Sigma-Aldrich) to verify successful transfer. Afterwards, membranes were blocked in 5 % milk (w/v) in TBS-T (200 mM Tris (Merck), 1.36 mM NaCl (Merck), 0.1 % v/v Tween 20 (Sigma)) for 1 h at room temperature. Primary antibodies were used in a 1:1,000 ratio in 5 % milk (w/v) in TBS-T for regular antibodies and 5 % BSA (w/v) in TBS-T for phospho-antibodies and incubated overnight at 4 °C (Supplementary Tab. 3). After washing 4x for 10 min with TBS-T, secondary antibodies were applied in a 1:10,000 ratio in 5 % milk (w/v) in TBS-T for 1 h at room temperature.

### RNA Sequencing (mouse)

For the mouse RNA-Seq, we isolated RNA from primary white adipocytes using the NucleoSpin® RNA kit as described above. We measured RNA concentration and quality by Nanodrop and shipped the samples to Novogene (UK), where they performed bulk mRNA-Seq with the Illumina Novaseq platform. Briefly, after quality control of the RNA samples, mRNA was enriched, fragmented and reverse-transcribed into cDNA. After library preparation, the samples were sequenced and the resulting reads were filtered and aligned to the mouse genome (GRCm39/mm39) using HISAT2 v.2.0.5^49^. We performed differential gene expression analysis using R v.4.3.0 with the DESeq2 v.1.44.0^50^ and ashr v.2.2-63^51^ packages, together with the dplyr v.1.1.4^52^ package for data handling. Further analysis and visualization of results were performed using ggplot2 v.3.5.1^53^, annotables v.0.2.0^54^, clusterProfiler v.4.12.0^55,56^, ComplexHeatmap v.2.20.0^57,58^, EnhancedVolcano v.1.22.0^59^ and VennDiagram v.1.7.3^60^ packages. Differentially expressed genes were filtered by *P*_adj_<0.05 and absolute log_2_ fold change >0.3 for comparing the effect of genetic manipulation in untreated cells and absolute log_2_ fold change >1 for cells treated with cholesterol for downstream analysis. For the human RNA-Seq data, RNA extraction from adipose tissue was performed using the SMARTseq^61,62^ protocol. Briefly, RNA was enriched and reverse transcribed using oligo(dT) and TSO primers, followed by cDNA amplification using *in silico* validated PCR primers. cDNA was processed using a Nextera DNA Flex Kit with Tn5 transposase. Single-end sequencing of all libraries took place on a Novaseq 6000 instrument at the Functional Genomics Center in Zurich. Raw sequencing reads underwent adapter and quality trimming using Fastp v0.20.0^63^, with a minimum read length of 18 nucleotides and a quality cut-off of 20. Read alignment to the human reference genome (assembly GRCh38.p13, GENCODE release 32) and gene-level expression quantification were performed using Kallisto v0.48^64^. Samples with read counts exceeding 20 million were down-sampled to 20 million read counts using the R package ezRun v3.14.1 (https://github.com/uzh/ezRun, accessed on 23 March 2022). The variance stabilizing transformation from DESeq2 v1.32.0^50^ was utilized for homoscedastic normalization based on library size. Batch effects were assessed using the R swamp package v1.5.1 (https://CRAN.R-project.org/package=swamp, accessed on 23 March 2022).

### Statistics

Data are expressed as the mean±standard error of the mean (s.e.m.). For comparing 2 groups, we used Student’s t test, when samples followed normal distribution or Mann-Whitney U-test where appropriate. We used one-way analysis of variance (ANOVA) for comparing three and more groups and two-way ANOVA for comparing two factors with post-hoc multiple comparisons. P values were adjusted for multiple testing where appropriate as indicated in the figure legends. Analyses were performed using R (4.3.0), Microsoft Excel and/or GraphPad Prism (10.0.1) and ImageJ. *P*<0.05 and *P*_adj_/q<0.05 were considered significant, as indicated by asterisks and letters in the figure legends.

## Author contributions

C.J. designed and conducted experiments, analyzed data, and wrote the manuscript. A.H., M.K., A.G., C.W., and M.B. analyzed human data. S.Ko., J.C., A.W., C.S., and S.Kh., B.A.A., and O.T.B performed experiments. Y.D., S.H., C.W., S.B.W, and G.S.H. designed experiments. A.B. designed and performed experiments, analyzed data, wrote the manuscript, and is the guarantor of the work. All authors read and commented on the manuscript.

## Acknowledgements

We thank Daniela Hass, Yvonne Jansen, Thomas Pitsch, and Silvia Weidner for excellent technical assistance. We thank Ana Paula Arruda, Mary Ebrahim, Kosei Eguchi, Karen Inouye, Henrika Jodeleit, Truc B. Nguyen, Kacey J. Prentice, Nicole A. Snyder, and Jonas Standt for their help with animal work. We thank the members of the Bartelt Lab for discussions and the enjoyable atmosphere and thank the members of the Hotamışlıgil Lab for stimulating discussions when this project was started. We thank Wenfei Sun and Hua Dong for supporting the RNA-Seq analysis. O.T.B. was supported by Deutsche Forschungsgemeinschaft (DFG, BR 5355/2-1), German Federal Ministry of Education and Research (BetterView), National Center for Tumor Diseases (SATISFIES), Helmholtz Imaging Project grant ZT-I-PF-4-038, Chan Zuckerberg Initiative Deep Tissue Imaging grants DTI0000000248 and DTI2-0000000206. M.B. received funding from grants from the DFG (209933838, SFB1052-B1) and by Deutsches Zentrum für Diabetesforschung (82DZD00601). A.B. was supported by the DFG (SFB1123-B10), the Deutsches Zentrum für Herz-Kreislauf-Forschung, and the European Research Council Starting Grant PROTEOFIT. We used BioRender.com for graphics. We apologize to colleagues whose work we could not cite due to space limitations.

## Data Availability

All data necessary to evaluate the conclusions in the paper are included in the paper. The human RNA-Seq data from the LOBB have not been deposited in a public repository due to restrictions imposed by patient consent but can be obtained from M.B. upon request.

## Supplemental material

**Supplementary Fig. 1:**
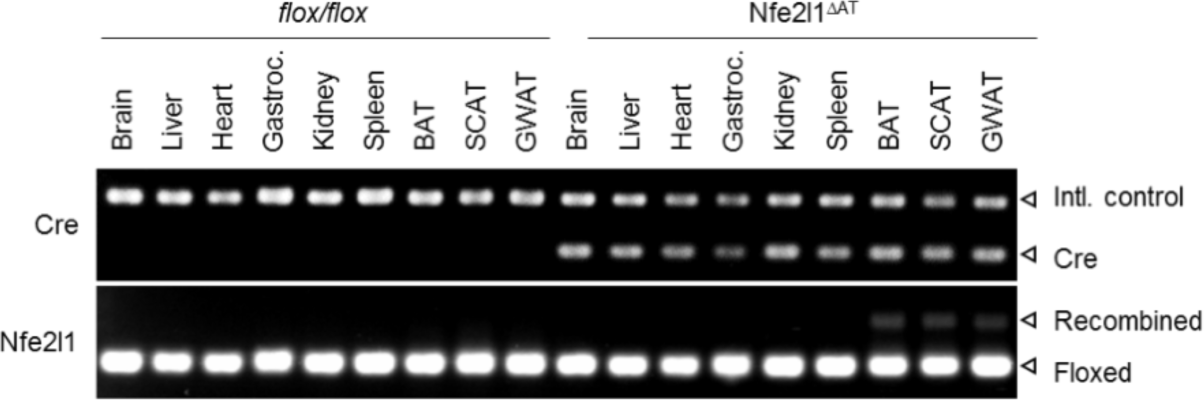
Adipose tissue-specific Cre-mediated recombination of the *Nfe2l1* gene. Intl.: Internal, Gastroc.: *Musculus Gastrocnemius*

**Supplementary Fig. 2:**
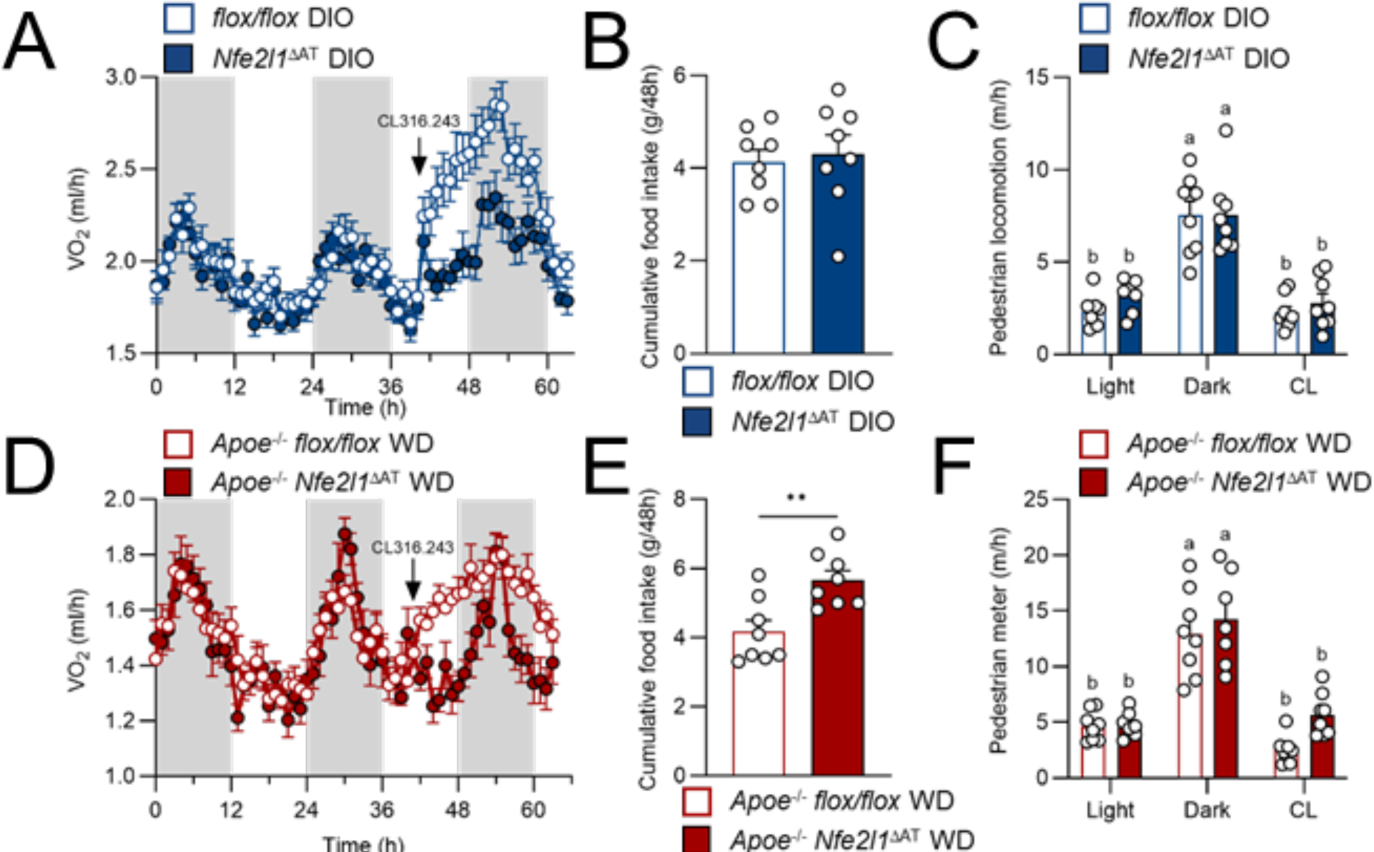
Energy metabolism of DIO *Nfe2l1*^ΔAT^ mice and WD *Apoe*^−/−^ *Nfe2l1*^ΔAT^. (A) Oxygen consumption rate (VO_2_) of *flox/flox* and *Nfe2l1*^ΔAT^ mice on DIO, n=8, data are mean+s.e.m. (B) Cumulative food intake over 48 h *flox/flox* and *Nfe2l1*^ΔAT^ mice on DIO, n=8, data are mean+s.e.m. (C) Activity of *flox/flox* and *Nfe2l1*^ΔAT^ mice on DIO, n=8, compact letter display by 2-way ANOVA with Tukey post-hoc test. (D) VO_2_ of *Apoe*^−/−^ *flox/flox* and *Apoe*^−/−^ *Nfe2l1*^ΔAT^ mice on WD, n = 7-8, data are mean+s.e.m. (E) Cumulative food intake over 48 h, n=7-8, data are mean+s.e.m., ** *P* < 0.01 by Student‘s T-test. (F) Activity, n=7-8, data are mean+s.e.m., compact letter display by 2-way ANOVA with Tukey post-hoc test.

**Supplementary Fig.3:**
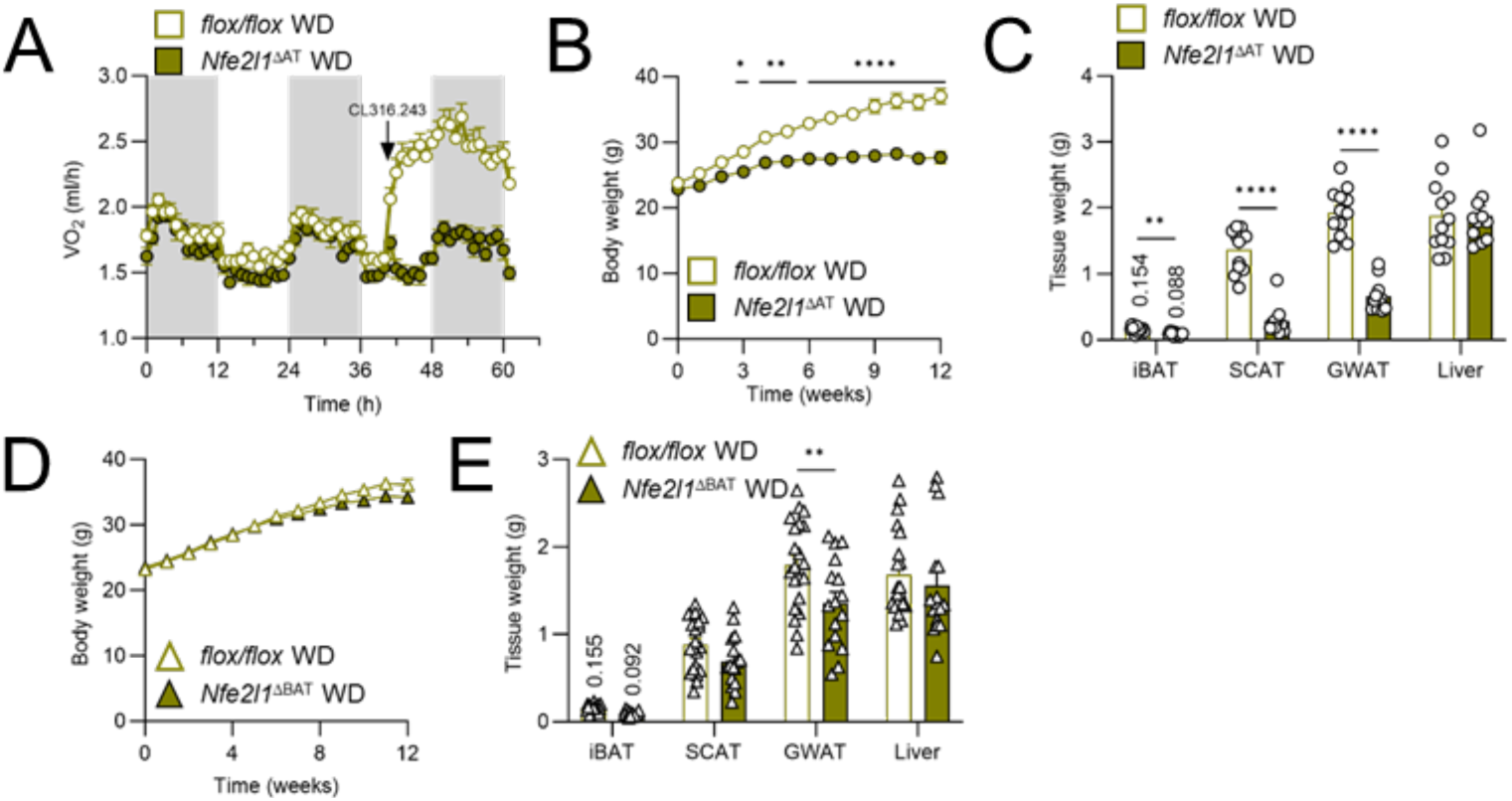
Adipose tissue phenotype of *Nfe2l1*^ΔAT^ and *Nfe2l1*^ΔBAT^ mice on WD. (A) Oxygen consumption rate (VO_2_) of *flox/flox* and *Nfe2l1*^ΔAT^ mice on WD, n=10-11, data are mean+s.e.m. (B) Body weight of *flox/flox* and *Nfe2l1*^ΔAT^ mice on WD, n=11-12, data are mean+s.e.m. (C) Tissue weights of *flox/flox* and *Nfe2l1*^ΔAT^ mice on WD, n=11-12, data are mean+s.e.m., \*\**P*<0.01, \*\*\*\**P*<0.001 by Student‘s T-test. (D) Body weight of *flox/flox* and *Nfe2l1*^ΔBAT^ mice on WD, n=16-21, data are mean+s.e.m. (E) Tissue weights of *flox/flox* and *Nfe2l1*^ΔBAT^ mice on WD, n=16-21, data are mean+s.e.m., \*\**P*<0.01, \*\*\*\**P*<0.001 by Student‘s T-test.

**Supplementary Figure 4.**
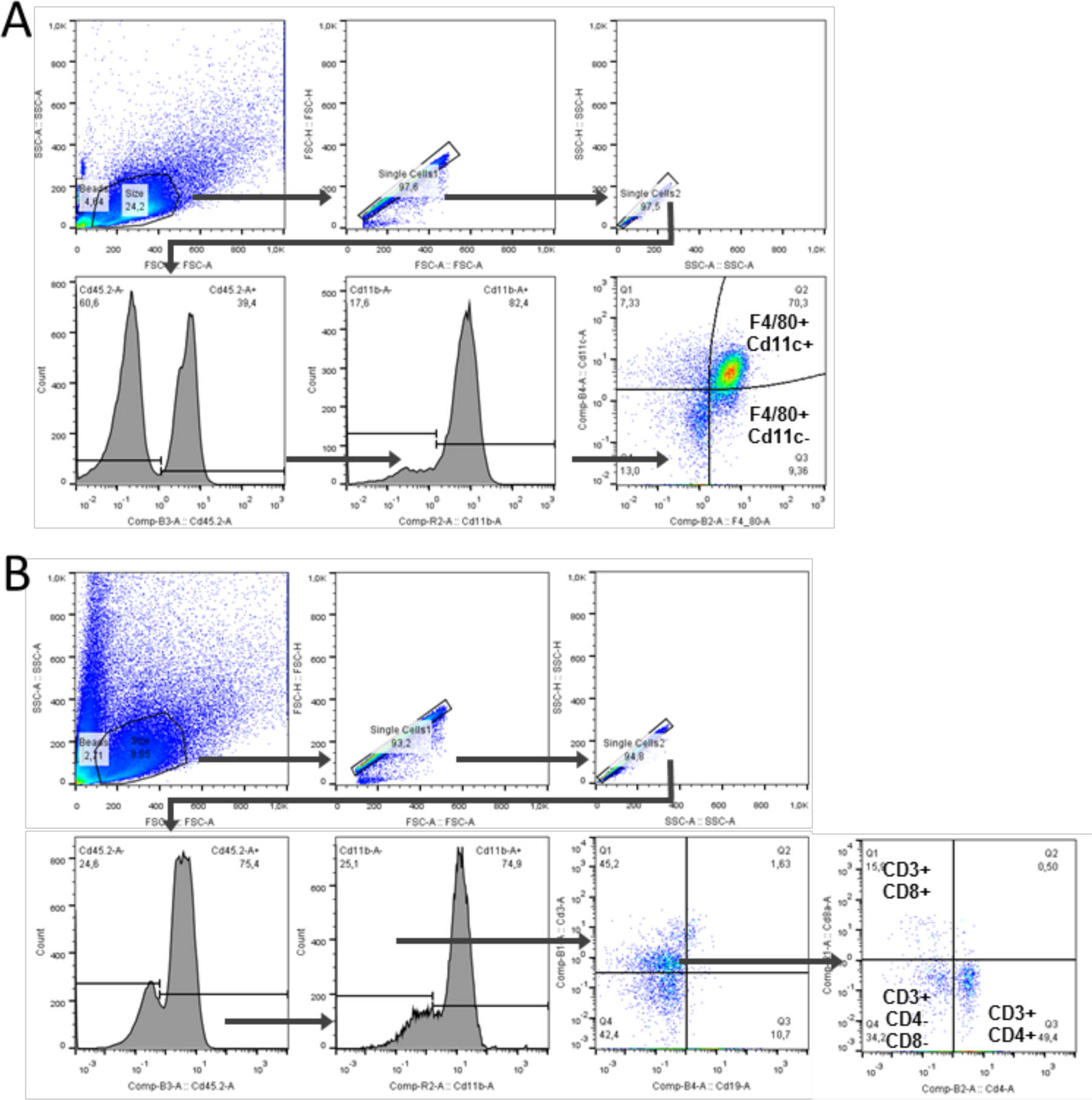
Flow cytometry gating strategies. (A) Gating strategy for F4/80+ cells. (B) Gating strategy for CD3+ cells.

**Supplementary Tab. 1:** FACS Antibodies.

| Antibody | Supplier | Order number | Working dilution |
| --- | --- | --- | --- |
| Purified anti-mouse CD16/32 Antibody (Fc Block) | BioLegend | 101302 | 1:100 |
| F4/80 Monoclonal Antibody PE | eBioscience | 12-4801-82 | 1:100 |
| Cd11b Monoclonal Antibody APC-eFluor 780 | eBioscience | 47-0112-82 | 1:125 |
| Cd45.2 Monoclonal Antibody PerCP-Cyanine5.5 | eBioscience | 45-0454-82 | 1:100 |
| Cd11c Monoclonal Antibody PE-Cyanine7 | BioLegend | 117318 | 1:100 |
| Cd4 Monoclonal Antibody PE | eBioscience | 12-0041-82 | 1:200 |
| Cd8a Monoclonal Antibody APC | BioLegend | 100712 | 1:200 |
| PE/Cy7 anti-mouse CD19 Antibody | BioLegend | 115519 | 1:800 |
| CD3e Monoclonal Antibody (145-2C11), FITC | eBioscience | 11-0031-85 | 1:100 |

**Supplementary Tab. 2:**
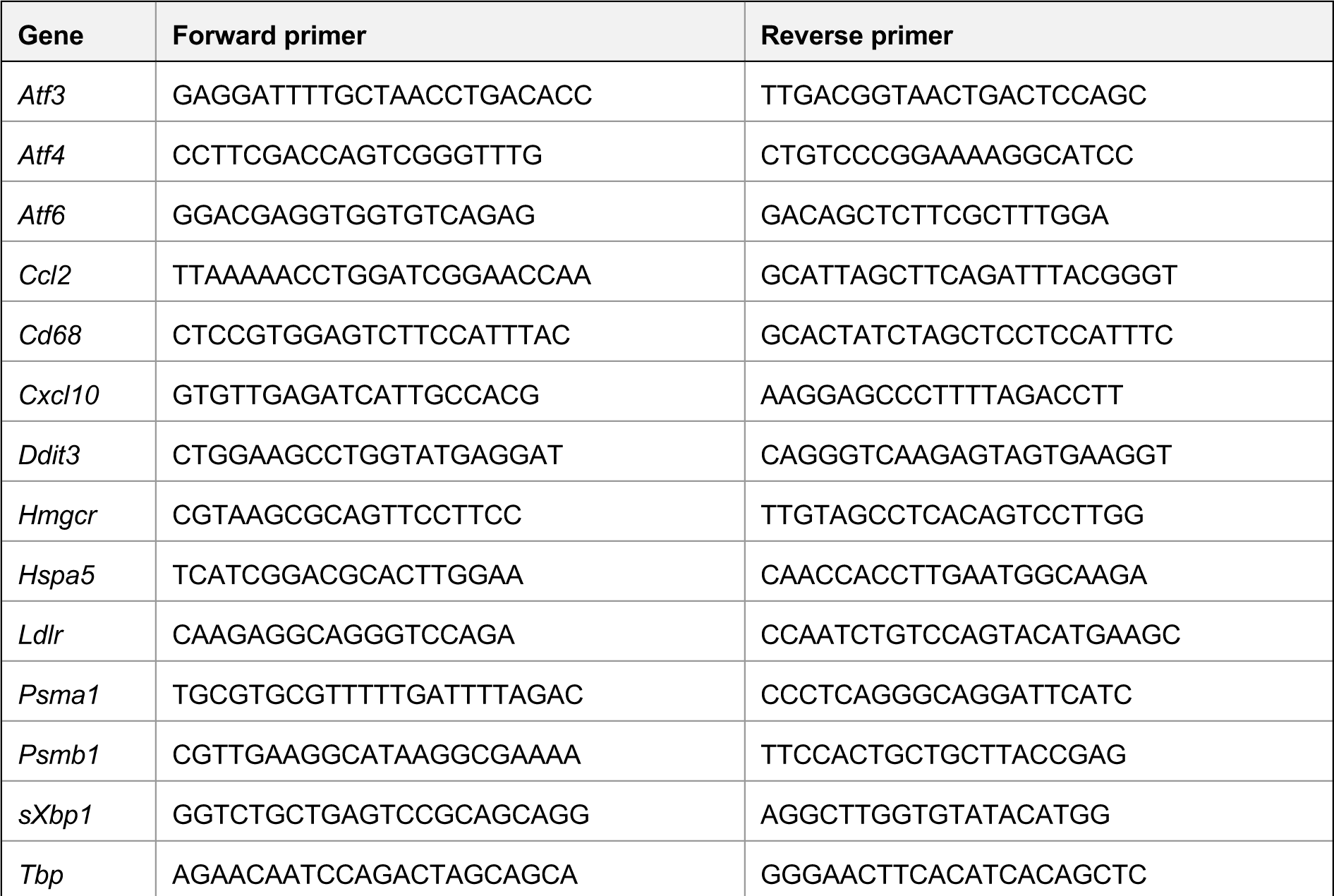
qPCR primers.

**Supplementary Tab. 3:**
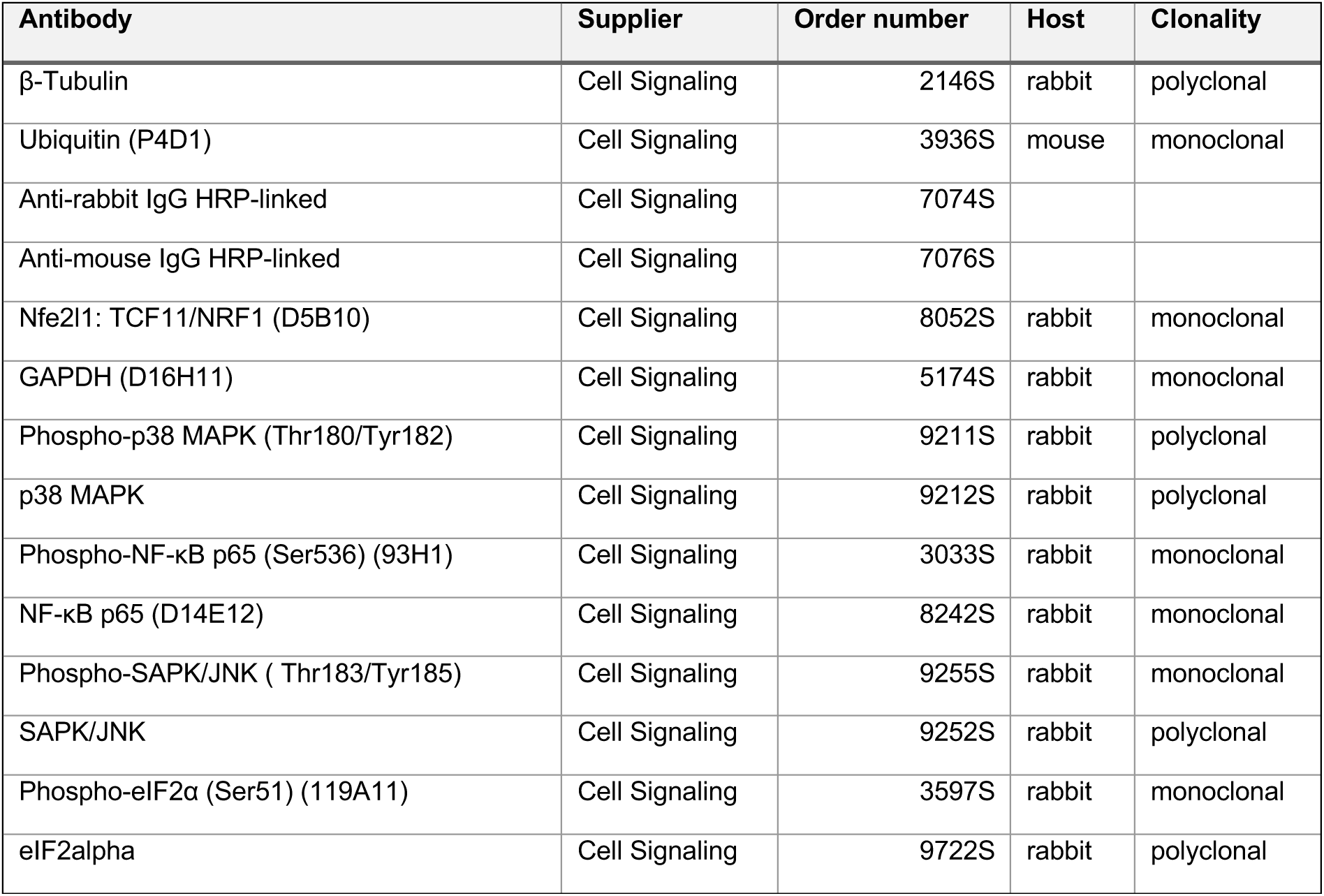
Immunoblots antibodies:

| Antibody | Supplier | Order number | Host | Clonality |
| --- | --- | --- | --- | --- |
| β-Tubulin | Cell Signaling | 2146S | rabbit | polyclonal |
| Ubiquitin (P4D1) | Cell Signaling | 3936S | mouse | monoclonal |
| Anti-rabbit IgG HRP-linked | Cell Signaling | 7074S |  |  |
| Anti-mouse IgG HRP-linked | Cell Signaling | 7076S |  |  |
| Nfe2l1: TCF11/NRF1 (D5B10) | Cell Signaling | 8052S | rabbit | monoclonal |
| GAPDH (D16H11) | Cell Signaling | 5174S | rabbit | monoclonal |
| Phospho-p38 MAPK (Thr180/Tyr182) | Cell Signaling | 9211S | rabbit | polyclonal |
| p38 MAPK | Cell Signaling | 9212S | rabbit | polyclonal |
| Phospho-NF-κB p65 (Ser536) (93H1) | Cell Signaling | 3033S | rabbit | monoclonal |
| NF-κB p65 (D14E12) | Cell Signaling | 8242S | rabbit | monoclonal |
| Phospho-SAPK/JNK ( Thr183/Tyr185) | Cell Signaling | 9255S | rabbit | monoclonal |
| SAPK/JNK | Cell Signaling | 9252S | rabbit | polyclonal |
| Phospho-eIF2α (Ser51) (119A11) | Cell Signaling | 3597S | rabbit | monoclonal |
| eIF2α | Cell Signaling | 9722S | rabbit | polyclonal |

